# Maintenance of species differences in closely related tetraploid parasitic *Euphrasia* (Orobanchaceae) on an isolated island

**DOI:** 10.1101/2020.04.29.067579

**Authors:** Hannes Becher, Max R. Brown, Gavin Powell, Chris Metherell, Nick J. Riddiford, Alex D. Twyford

## Abstract

Polyploidy is pervasive in angiosperm evolution and plays important roles in adaptation and speciation. However, polyploid groups are understudied due to complex sequence homology, challenging genome assembly, and taxonomic complexity. Here we study adaptive divergence in taxonomically complex eyebrights (*Euphrasia*), where recent divergence, phenotypic plasticity and hybridisation blur species boundaries. We focus on three closely-related tetraploid species with contrasting ecological preferences, and which are sympatric on Fair Isle, a small isolated island in the British Isles. Using a common garden experiment, we show a genetic component to the morphological differences present between these species. Using whole genome sequencing and a novel *k*-mer approach, we demonstrate an allopolyploid origin, with sub-genome divergence of approximately 5%. Using ~2 million SNPs we show sub-genome homology across species consistent with a common origin, with very low sequence divergence characteristic of recent speciation. This genetic variation is broadly structured by species, with clear divergence of Fair Isle heathland *E. micrantha,* while grassland *E. arctica* and coastal *E. foulaensis* are more closely related. Overall, we show tetraploid *Euphrasia* is an allopolyploid system characterised by postglacial species divergence, where adaptation to novel environments may be conferred by old variants rearranged into new genetic lineages.

## Introduction

Plant populations that grow in contrasting ecological conditions will experience different selection pressures for adaptive traits that underlie survival and reproduction (Clausen et al., 1940; Clausen et al., 1948; Clausen and Hiesey, 1958; Núñez-Farfán and Schlichting, 2001). This divergent ecological selection may cause adaptive divergence of populations, and lead to the origin of novel ecotypes and species (McNeilly and Antonovics, 1968; Böhle et al., 1996; Baldwin and Sanderson, 1998; Nevado et al., 2016; Favre et al., 2017). The trajectory of divergence in the early stages of speciation is complex, with recent studies showing that populations may diverge in the face of ongoing gene flow that was previously thought sufficient to homogenise population differences and oppose divergence (Danley et al., 2000; Papadopulos et al., 2011; Nadeau et al., 2013; Richards et al., 2016). While such insights have been made in different plant species, these are mostly ecological and evolutionary model systems that are amenable to genomic analysis (Bernasconi et al. 2009, Twyford et al. 2015). There are numerous plant groups that are under-represented in current speciation genomic studies, and these include species that are characterised by recent polyploidy and groups with complex taxonomy where species boundaries are poorly understood.

Polyploidy, or whole genome duplication (WGD), is common in angiosperms, with all extant species having experienced at least one round of polyploidy (Wendel et al., 2016). WGD is frequently linked to adaptation and speciation in plants (Wood et al., 2009; Levin, 2019). It may facilitate adaptation on its own in autopolyploids (Baduel et al., 2019), which often show different ecological preferences to their diploid progenitors (Parisod et al., 2010). In conjunction with hybridisation, WGD creates allopolyploids, where redundant genomic variants can take on novel fates (neofunctionalisation) or partition current functions (subfunctionalisation) (Cheng et al. 2018). Polyploidy increases genome size, which itself may affect fitness (Guignard et al., 2016), and it can increase short-term adaptive potential allowing organisms to colonise new environments and to tolerate stressful conditions (Lowe and Abbott, 1996; Ainouche et al., 2009; Soltis et al., 2012).

While polyploidy is now widely appreciated as a key driver of plant diversification (Tank et al., 2015; Ren et al., 2018), there are considerable challenges in the study of polyploidy that limit our understanding of this key evolutionary process. First, polyploidy is common in taxonomically complex groups and certain apomictic taxa, where species limits may be uncertain and where taxon identity is unknown (Popp et al., 2005; Guggisberg et al., 2006; Brysting et al., 2007). Second, comparative genomics of polyploids relies on correctly determining sequence homology, which is challenging with the additional gene copies from genome duplication (homeologs). Third, reference genome assembly, which is critical for many aspects of speciation genomics, such as genome scans for detecting outlier regions subject to selection (Ravinet et al., 2017; Campbell et al., 2018) is notoriously difficult in polyploids (but not impossible, Hollister et al., 2012). One way of circumventing the inference of sequence homology in a polyploid genome assembly is the use of methods based on DNA *k*-mers (DNA ‘words’ of fixed length *k*). Such approaches have recently been used to infer ploidy and heterozygosity in polyploid samples, and to identify genomic regions associated with a phenotype (Mapleson et al., 2016; Ranallo-Benavidez et al., 2020; Voichek and Weigel, 2020). They could be extended to further characterise polyploid genome structure based on a demographic model of population divergence, without the need for a reference genome (Lohse et al., 2011; Lohse et al., 2016).

The arctic and boreal regions of northern Europe are renowned for their diversity of polyploid taxa (Stebbins, 1984; Abbott and Brochmann, 2003; Brochmann et al., 2004), with eyebrights *(Euphrasia),* a genus of ~263 species (Twyford, Unpublished), one of the most diverse. *Euphrasia* species are infamous for their complex morphological diversity, with many forms grading into others and being relatively indistinct. This taxonomic complexity in *Euphrasia* is a consequence of a diverse set of factors; the genus is characterised by recent rapid postglacial divergence, with species hybridizing extensively (Yeo, 1966; Stace, 2019). There is also considerable variation in ploidy, and some species are highly selfing (Yeo, 1966; French et al., 2008; Stone, 2013). Moreover, *Euphrasia* are generalist facultative hemiparasites that are green and photosynthesise, but also attach to a host plant and steal nutrients and water. The growth of *Euphrasia* depends on the host species, with this phenotypic plasticity further contributing to taxonomic complexity (Brown et al., 2020). While an incredibly complex genus, genomic studies of *Euphrasia* promise to reveal key insights into the nature of species differences and how hybridisation, selfing and phenotypic plasticity shape the evolution of polyploid taxa.

Given the scale of taxonomic complexity in *Euphrasia,* we choose to focus on co-occurring species on a single island. Fair Isle, a small island of 768 hectares, is the most remote inhabited island in the British Isles, situated halfway between Orkney (42 km away) and mainland Shetland (39 km away). Fair Isle *Euphrasia* provide an ideal study system due to their isolation from nearby gene flow and the closely intermixed habitats that support different species, the presence of only tetraploid *Euphrasia,* and the well-characterised island flora of 260 UK native species and 71 aliens (Quinteros Peñafiel et al., 2017), including eight *Euphrasia* species and eight putative hybrids. Here we focus on three tetraploid *Euphrasia* species that are widely distributed across the island but differ in their habitat preferences (see Figure 1). These species are not distinguishable by ITS or plastid-based barcoding (Wang et al., 2018). *Euphrasia arctica* is a grassland species that has vigorous growth and large flowers that are assumed to be highly outcrossing. *Euphrasia foulaensis* is a coastal specialist with thickened leaves and compressed growth. It has small flowers, some of which do not open and are thought to be cleistogamous. *Euphrasia micrantha* is an upland heathland species that is slender and often has pink-suffused leaves, stems and flowers. Its very small flowers are highly selfing (Stone, 2013). *E. micrantha* is variable in morphology throughout its range, with the Fair Isle form smaller, usually unbranched, with magenta flowers with a less distinct lower floral ‘lip’. Despite the three species having different habitat preferences, the small scale of the island and fine-scale habitat differences means species occur in sympatry *(sensu* Mallet et al. 2009) (Figure 1). To place these Fair Isle plants in a broader context, we also include a sample of diploid and tetraploid individuals from elsewhere in Great Britain.

**Figure 1.**
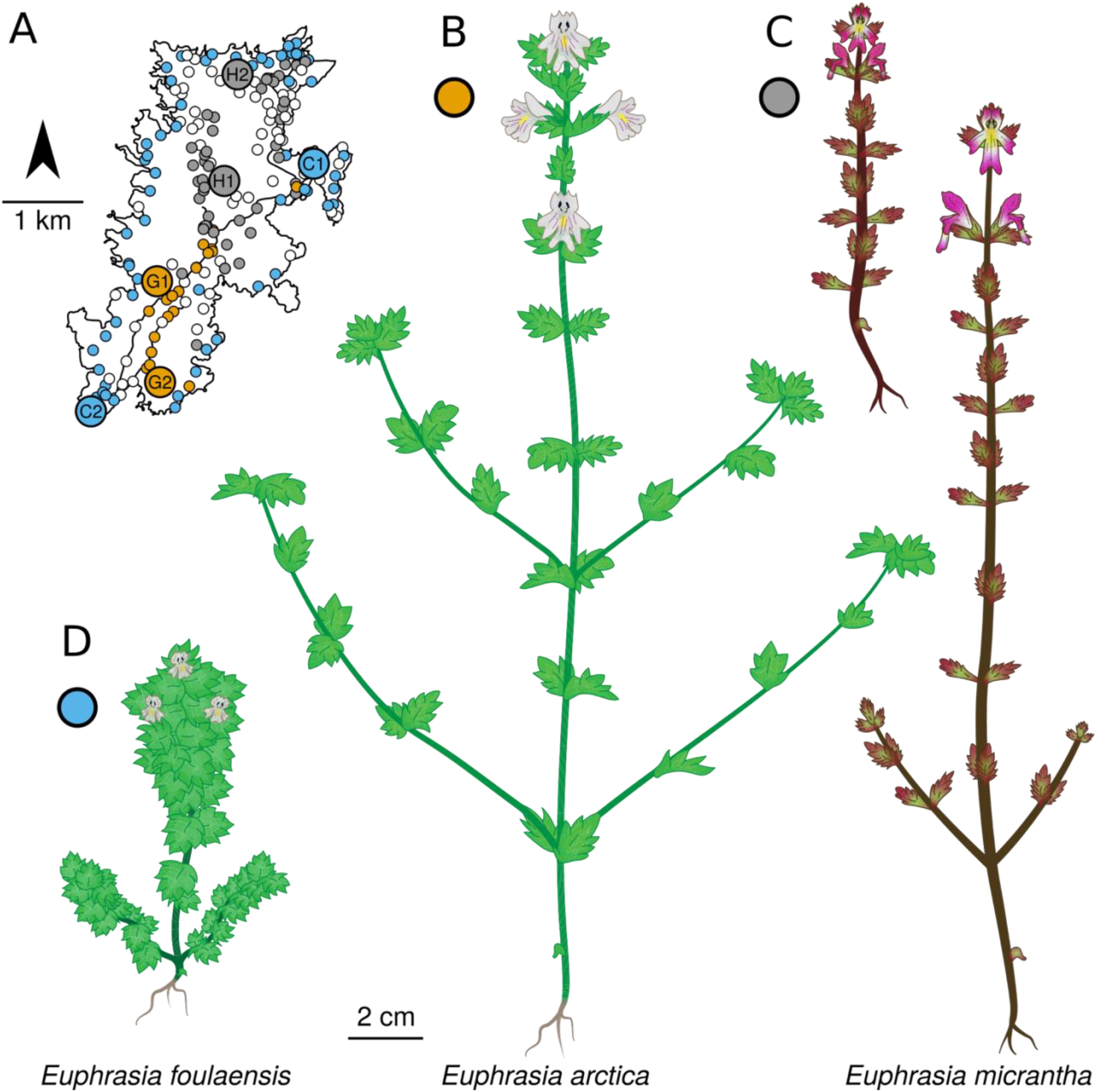
The three focal *Euphrasia* species and their distributions on Fair Isle. **A)** Geographic distributions of *E. arctica*, *E. foulaensis* and *E. micrantha* based on a field survey recording 282 presence (coloured) or absence (white) census points. Labels indicate the populations from which seeds were sourced and traits were measured. B) *E. arctica* C) *E. micrantha.* The smaller form is typical for Fair Isle. D) *E. foulaensis.*

In this study we use field-based observations, a common garden experimental approach, and whole genome sequencing to understand the nature of species differences in a group of taxonomically complex, sympatric, polyploid *Euphrasia* species. We address the following research questions: (1) is there a genetic component to the morphological differences between species with different habitat preferences?, (2) what is the evolutionary history of polyploidisation in *Euphrasia?,* and (3) what is the landscape of genomic differentiation between sympatric species? To address question 2, we propose a novel *k*-mer based analytical approach to infer sub-genome divergence in allopolyploids. Our results show how species differences are maintained over a fine-spatial scale despite incomplete reproductive isolating barriers and reveal how genomic approaches can be used to characterise speciation histories of a non-model tetraploid group.

## Results

### Phenotypic differentiation between *Euphrasia* species is heritable

To understand morphological differences among Fair Isle *Euphrasia* species, we compared trait variation in the wild, and in a common garden experiment. We confirmed that in the wild *E. arctica* is tall (mean height = 80.0 mm) and large flowered (mean corolla size = 7.6 mm); *E. foulaensis* short (19.7 mm) and smaller flowered (5.9 mm) and *E. micrantha* intermediate in height (40.2 mm) and small flowered (4.7 mm), with trait these values significantly different between all three species in mixed-effect models (p <0.01; Fig. 2A, Supplemental Table 1, Supplemental Table 2, and Supplemental Text 1). Overall however many more traits showed significant differences at the population rather than the species level (Fig. 2A, bottom right triangles). Based on multi-trait phenotypes, individuals partly cluster by species in a principal component analysis (PCA, Fig. 2B), while linear discriminant analysis (LDA) with cross-validation, classified 96% of individuals to the species we had assigned. These results show species and populations differ in their overall phenotype and in some key traits in natural populations.

**Figure 2.**
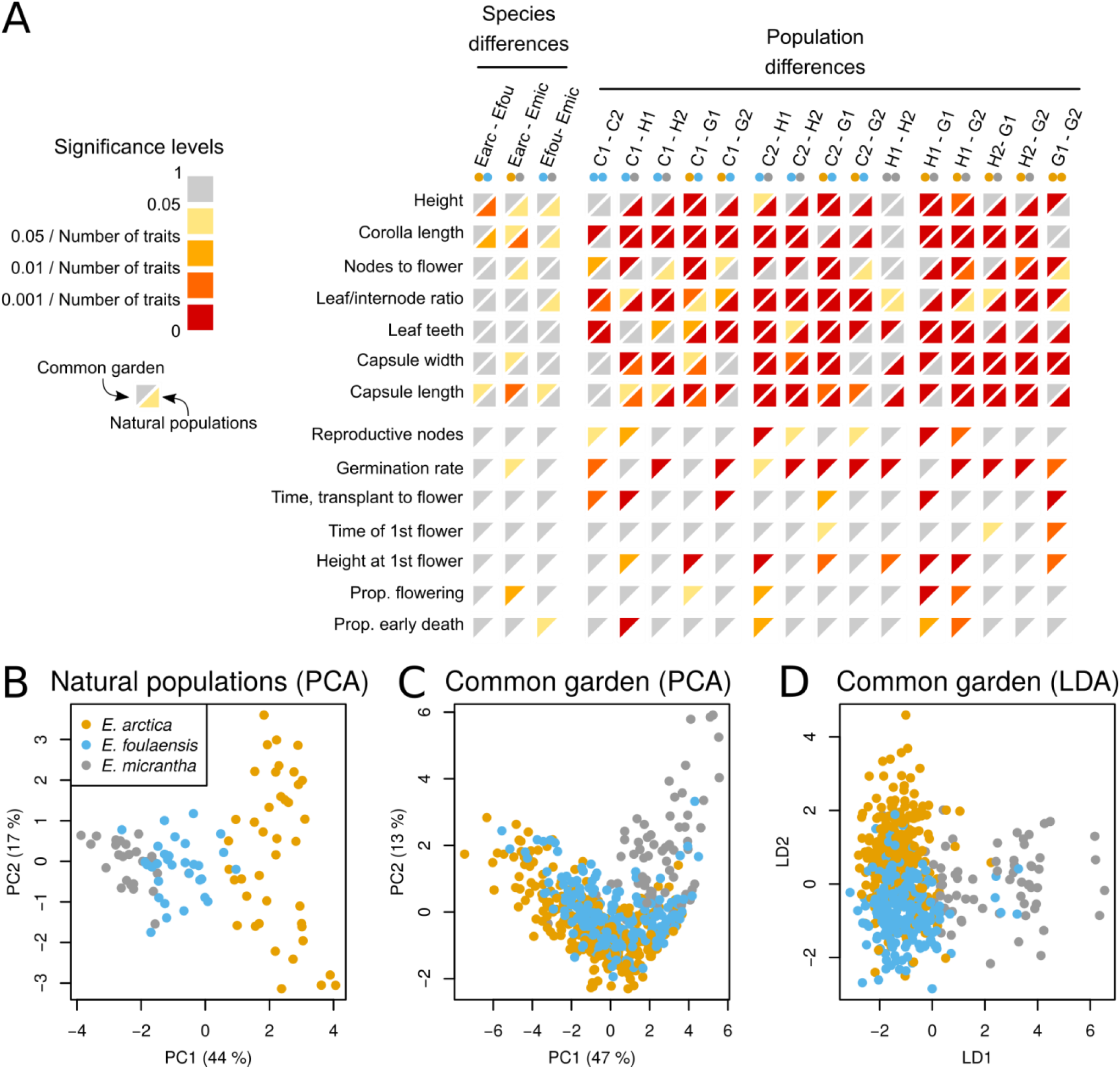
Morphological trait differentiation between three species of *Euphrasia* in a common garden experiment and in natural populations. A) Significance levels of trait differences between species (left) and populations (right) from field measurements in natural populations (bottom right triangles) and from the common garden (top left triangles, see Supplemental Text 1 for the magnitude of difference and Supplemental Table 2 for the means and standard errors associated). Comparisons within rows are corrected for multiple testing. Within columns, the colour scale is the p-value corrected for the number of traits tested (seven in natural populations and 14 in the common garden). While significant trait differences are rare between species, they are numerous between populations. B) A PCA of trait measurements from natural populations shows separate clusters per species. C) A PCA of trait measurements taken from plants grown in the common garden shows little grouping by species. D) An LDA separates species in the common garden.

To assess whether the phenotypic species differences have a genetic component, we analysed morphological differentiation in a common garden setting, measuring 2116 individuals grown from seeds sourced from the same natural populations as above. Because host identity can affect performance in *Euphrasia* (Brown et al., 2020), we grew each individual with one of twelve different host species, which occur on Fair Isle (Supplemental Table 2). Including host species as a random effect improved the fit of our models, which then had significantly lower AIC values. Under these common garden conditions, there were fewer observable trait differences than in natural populations. The only trait showing significant differentiation between all species was capsule length, which is high in *E. arctica* (5.8 mm), intermediate in *E. foulaensis* (4.9 mm), and short in *E. micrantha* (3.7 mm) (p<0.01). However, there were still some pairwise species differences, and many pairwise population differences (Fig. 2A, top-left triangles). Overall, clustering of different species based on multi-trait phenotypes in the common garden (Fig. 2C) was much less clear than in natural populations (Fig. 2B). However, using LDA with cross validation it was possible to accurately classify 75% of individuals (i.e. 412 out of 549 individuals without any missing data were assigned the species from which their seeds had been collected, Fig. 2D). The success of LDA classification was comparable across species, with *E. arctica* and *E. foulaensis* misclassified as the other species 25% and 20% of the time, respectively, while *E. micrantha* was most commonly misclassified as *E. foulaensis* (18%). To test whether this pattern could be observed by chance, we ran LDA with cross validation on 9999 data sets with randomly reassigned species labels. The 99%-confidence interval of the resulting distribution ranges from 25% to 40%, indicating that the success rate of 75% is highly significant and was not the result of over fitting. Overall, the presence of species-specific multi-trait combinations in a common environment shows there is a genetic component to the phenotypic species differences, but the lower classification success in a common garden indicates that many trait differences observed in the wild are due to plasticity.

### Complex patterns of plastid genome and rDNA relatedness

We generated whole genome sequencing data for 18 *Euphrasia* individuals. These included twelve Fair Isle tetraploids from our three focal species and two individuals considered putative hybrids (labelled X1 and X2). The other samples were two tetraploids and four diploids from mainland Britain, see Supplemental Table 3 for details. *De novo* assembly of plastid genomes revealed complex patterns of plastid haplotype sharing and relatedness. Plastid genomes were similar in size (144,739-145,009 bp) and in sequence (>99.8% pairwise sequence identity). Fair Isle *Euphrasia* fell into three broad plastid haplogroups in phylogenetic analysis (Figure 3A). Haplogroup one is composed of a mix of Fair Isle *Euphrasia* and mainland British diploid and tetraploid species. Haplogroup two is predominantly found in Fair Isle samples, plus the putative hybrid species *E. rivularis* sampled from England. Haplogroup three is exclusively composed of Fair Isle individuals of *E. micrantha.* Within each of these haplogroups there are variable levels of genetic divergence, with some extremely closely related haplotypes differing by a few SNPs, and some more divergent haplotypes showing structural genetic changes. Further, there is plastid haplotype sharing between co-occurring samples of *E. arctica* and *E. foulaensis* on a roadside in the south of Fair Isle, as well as their putative hybrid (sample X1). Overall, these results indicate that diverse plastid haplotypes are maintained within an island population of *Euphrasia,* with Fair Isle *E. micrantha* differing from intermixed *E. arctica* and *E. foulaensis*.

**Figure 3.**
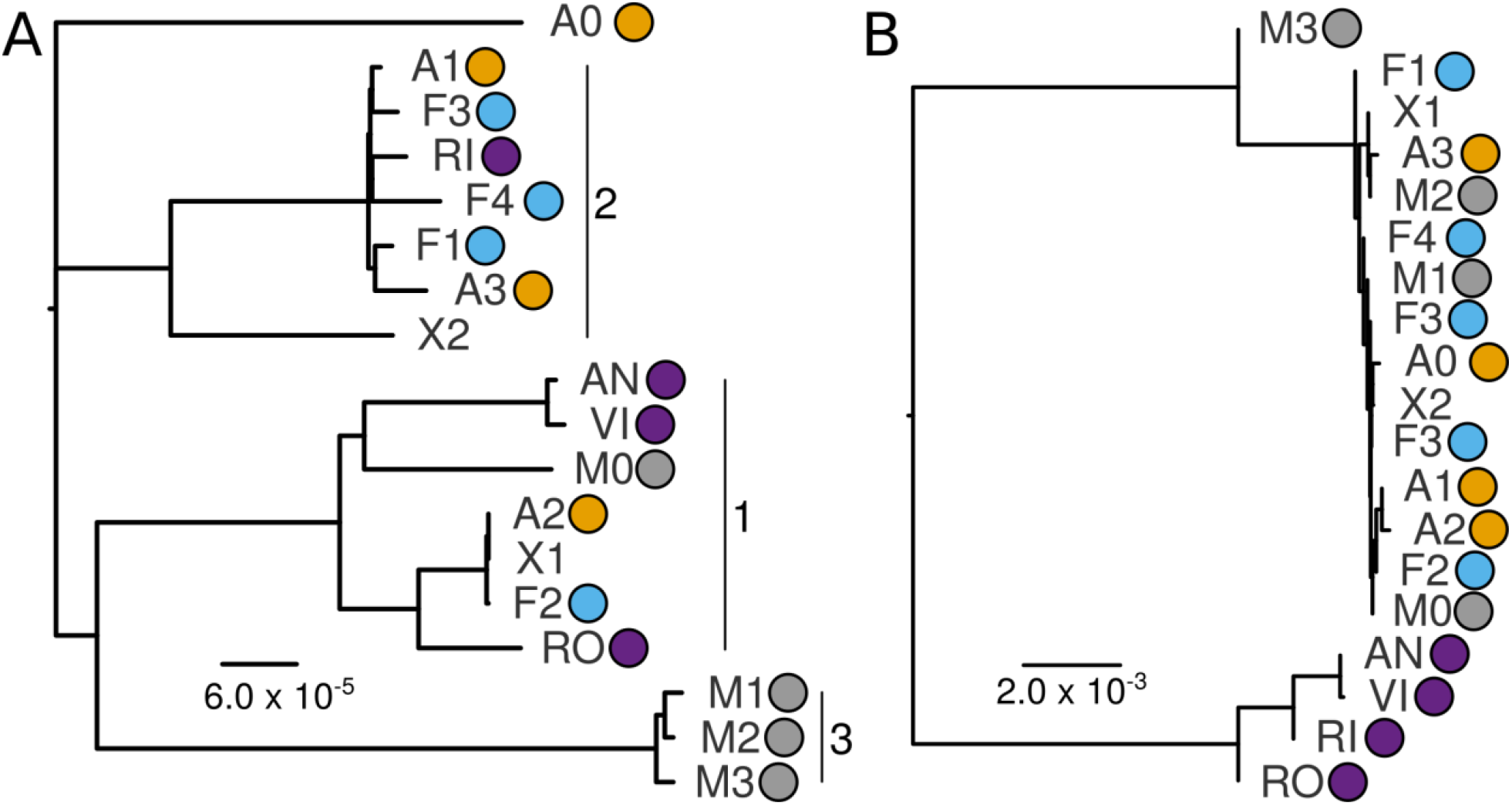
Evolutionary relationships of British *Euphrasia* plastid genomes rDNA sequences. A) Phylogenetic analysis of plastid genomes performed using a maximum-likelihood approach implemented in IQ-TREE with the K3Pu+F+I substitution model. B) Phylogenetic analysis of nuclear rDNA sequences using neighbour-joining approach implemented in Geneious 11. There were two rDNA haplotypes in sample F3. Coloured circles indicate species identity as in Figure 1, with diploids coloured in purple. Branch lengths indicated by the scale bar.

*De novo* assembly and comparative analyses of *Euphrasia* rDNA revealed deep divergence between UK diploids and tetraploids, confirming previous results from ITS sequencing of a broader taxonomic sample (Wang et al., 2018). Across the 5832bp rDNA coding there was 2.1% mean pairwise diploid-tetraploid divergence, but up to 10.8% divergence in the ITS2 region. Within tetraploids however, there was very limited sequence divergence, with >99.5% pairwise sequence identity between Fair Isle samples. rDNA haplotypes were shared between some individuals such as *E. foulaensis* F2 and *E. arctica* A1, with one sample of *E. micrantha* maintaining a more divergent haplotype. These results support recent divergence of species and populations, with extensive haplotype sharing particularly among *E. arctica* and *E. foulaensis*.

### A common allopolyploid origin with a shared diploid progenitor

*K*-mer spectra are widely used to assess the completeness of genome assemblies, to estimate genome sizes, and to estimate heterozygosity in diploids. To assess the genomic properties of tetraploid *Euphrasia* without the need for a genome assembly, we developed mathematical models for the shape of *k*-mer spectra of auto- and allotetraploids. We implemented these in a “shiny” App, Tetmer, which we then used on our data (see Figure 4A, Supplemental Text 2 and Supplemental Text 3). The *k*-mer spectra of all our tetraploids have a prominent 2x peak, containing more *k*-mers than all other peaks. Such spectra are not plausible under the autotetraploid model, unless implausible (negative) mutation rates are assumed, but can be explained by the allotetraploid model. Using Tetmer, we estimate the haploid *k*-mer number, which corresponds to the amount of single-copy sequence in a haploid genome, to be 185-225Mb in our samples. We then used Tetmer to estimate heterozygosity within and divergence between sub-genomes (see Figure 4B and C). The heterozygosity estimates are noisy because they depend mainly on the 1x peak in the *k*-mer spectra (Figure 4A), which is often partly concealed by sequencing errors and contaminations. We also estimated heterozygosity from SNPs called relative to a reference genome (described later), which showed similar patterns. The average *k*-mer based heterozygosity estimate over all samples is 0.2% and does not differ significantly between diploids and tetraploids (ANOVA, p_df=16_=0.07). Two samples, RO (*E. rostkoviana*) and A3 (*E. arctica*), have heterozygosity values that were considerably higher (1.1% and 0.52%), which likely corresponds to recent outcrossing events in these mixed-maters.

**Figure 4.**
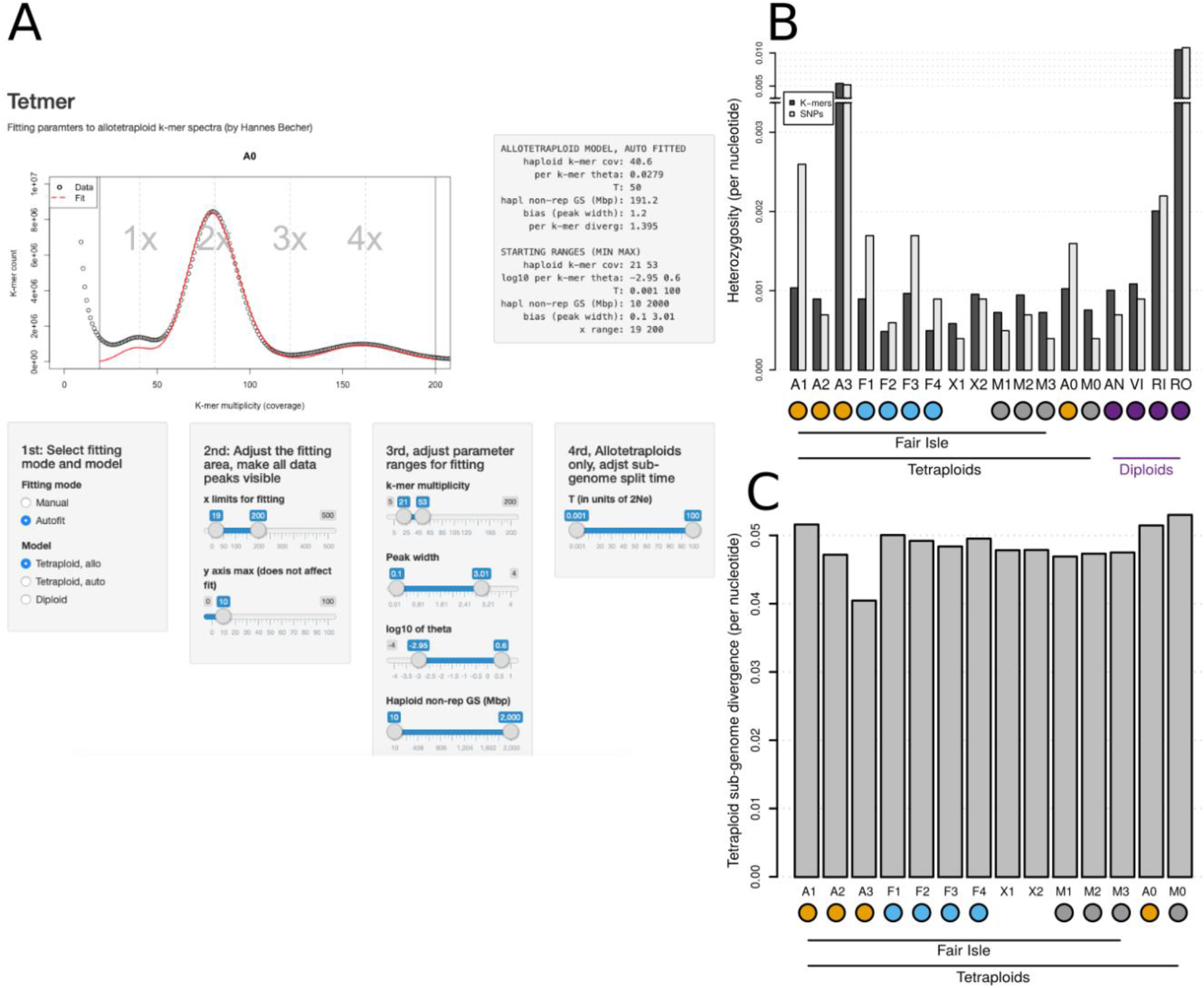
Estimates of heterozygosity and sub-genome divergence in allotetraploids based on *k*-mer spectra. A) A screenshot of our app “Tetmer”, which allows heterozygosity to be estimated within and nucleotide divergence between the sub-genomes of allotetraploids (it can also estimate heterozygosity in diploids and autotetraploids). B) Estimates of heterozygosity of *Euphrasia* individuals based on *k*-mers (dark bars) and SNPs (light bars). C) Estimates of the sub-genome divergence for all tetraploid *Euphrasia* individuals.

In Tetmer, we modelled the divergence between allotetraploid sub-genomes in an analogous way to diverging populations (see Methods and Supplemental Text 2). The longer sub-genomes have evolved independently the more fixed private alleles they have, which will increase the size of a *k*-mer spectrum’s 2x peak. Because the divergence estimate depends mainly on the relative size of the 2x and 4x peaks, it will be more accurate than heterozygosity estimates based on the 1x peak. Using Tetmer, we found that all tetraploids show a per-nucleotide sub-genome divergence of approximately 5%, one to two orders of magnitude higher than the heterozygosities observed (Figure 4C). The consistency of sub-genome divergence across samples raises the possibility of a common origin to these polyploids, involving similar parental progenitors.

To further our understanding of sub-genome divergence and polyploid history, we generated a draft genome assembly of a geographically isolated tetraploid sample of *E. arctica* from North Berwick, Scotland (individual A0). This hybrid assembly from 97x short read Illumina and 16.7x long-read Pacific Biosciences (PacBio) data comprises 1,009,737 scaffolds and spans 694Mbp in length. While fragmented, the assembly is relatively complete, as indicated by the KAT completeness plot (Supplemental Figure 1) and the BUSCO completeness score of 81.7%. We then mapped genome resequencing data of all 18 diploids and tetraploid individuals to our reference assembly and classified scaffolds based on mapping depths. Only 0.1% of scaffolds (1024 scaffolds; 3Mb) had 4x mapping depth in the tetraploids, as would be expected if they were autotetraploids, providing further support for an allopolyploid origin of the tetraploids. In contrast, 10,644 scaffolds (132Mb) had a diploid-level (2x) mapping depth across all tetraploids, representing regions where diverged (homeologous) sequences are assembled into separate scaffolds, which we call the “tetraploid” set. A subset of the tetraploid scaffolds had 2x mapping depth across all diploids and tetraploids and represent homologous regions across all our samples, the other scaffolds are mostly missing from diploids. We call these 3,454 scaffolds (46Mbp) the “conserved” set. The tetraploid and conserved sets form the basis for subsequent population genomic analyses (Figure 5A). Overall, the consistency in mapping depth patterns between tetraploids suggests widespread sequence homology and a common origin. Diploid-tetraploid homology supports the prediction that a (relative of a) British diploid was a progenitor of British tetraploids (Yeo, 1966), with a second sub-genome contributed by a divergent taxon.

**Figure 5.**
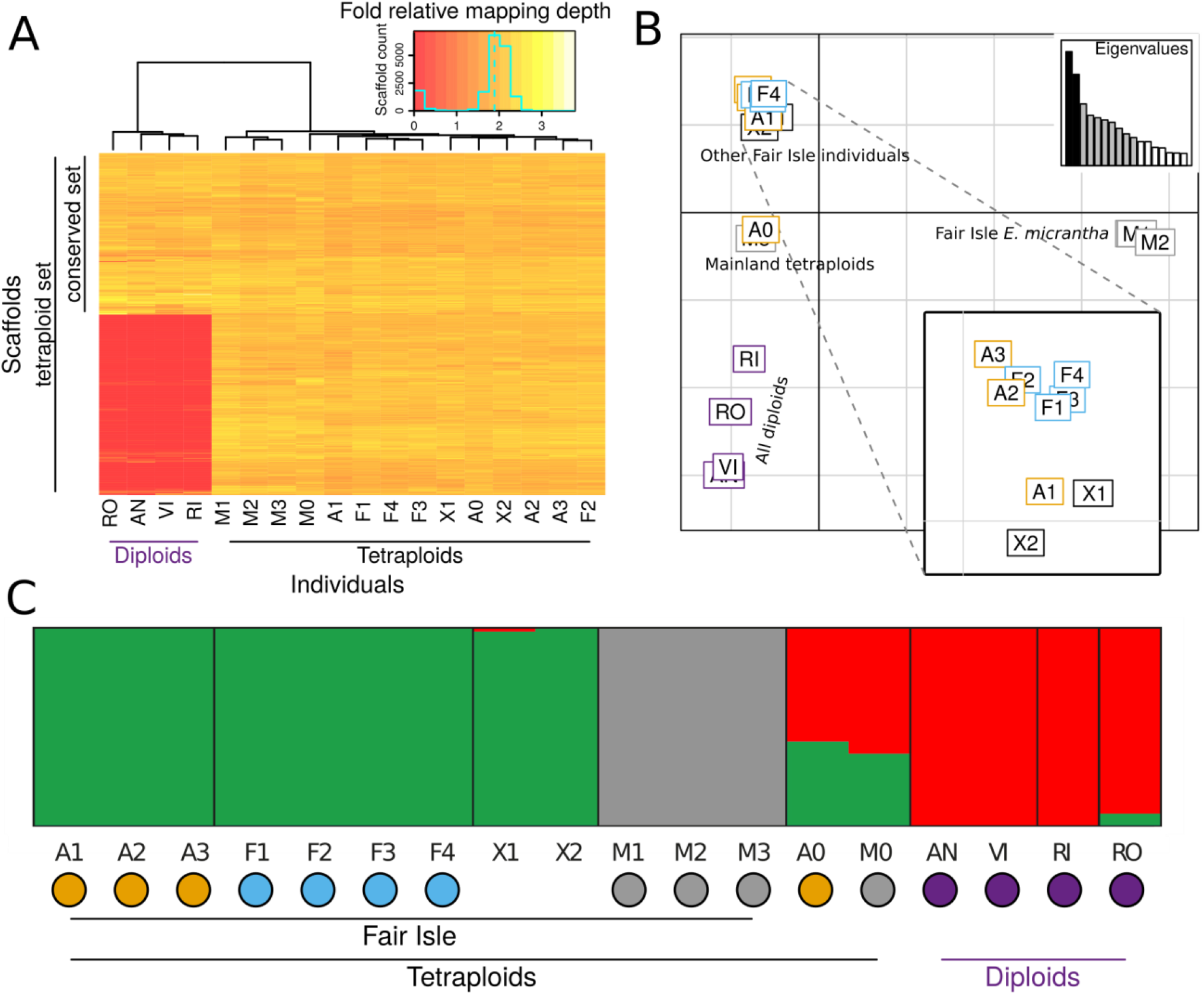
Diploid-tetraploid scaffold homology and the clustering of *Euphrasia* populations. A) Relative mapping depth in the tetraploid (2x depth in tetraploids) and conserved (2x depth all individuals) scaffold sets. Colours represent mapping coverage (see insert), tetraploid scaffolds not contained in the conserved set have low mapping depths in diploids indicative of absence (coloured red). B) PCA of genomic data separates Fair Isle *E. micrantha* individuals from other *Euphrasia.* PC2 separates tetraploids and diploids. Analysis based on 3454 SNPs with one SNP per scaffold. C) STRUCTURE analysis shows Fair Isle *E. micrantha* as a separate genetic cluster. Analysis based on the same SNP data as B. The plot shows the result for three genetic clusters (*K*=3), with individuals on the x-axis and admixture proportions on the y-axis. The mainland tetraploids, M0 and A0, are inferred to be admixed. Coloured dots in panels B and C represent taxa, to match Figure 3.

### Strong genetic structure despite low divergence

We used genetic clustering approaches to understand whether species from contrasting habitats are genetically cohesive. STRUCTURE analyses revealed that Fair Isle *E. micrantha* is a distinct genetic cluster in our sample of natural *Euphrasia* populations. Assuming three genetic clusters (*K*=3, Figure 4C), there are clusters corresponding to Fair Isle *E. micrantha,* other Fair Isle taxa, and diploids. The mainland tetraploids (A0 and M0) are found to be admixed between clusters. At higher *K*-values (Supplemental Figure 2), genetic clustering corresponds to broad taxonomic groupings, though *E. arctica* and *E. foulaensis* on Fair Isle are not separated. In a PCA, Fair Isle *E. micrantha* are separated from all other species on principal component (PC) 1, while other Fair Isle species are separated from diploids on PC2, with mainland tetraploids in between these groupings. Overall, these analyses point to divergence between Fair Isle *E. micrantha* and all other taxa being the major axis of divergence among our samples, rather than diploid-tetraploid divergence as found in other genetic analyses (French et al., 2008; Wang et al., 2018).

To better understand how variation is maintained within and between populations and species, we characterised genomic diversity and divergence based on 922,927 high-quality biallelic SNPs in the conserved and 2,166,914 SNPs in the tetraploid scaffolds. The per-species estimates of nucleotide diversity (H_e_) are considerably higher than individual heterozygosities, ranging from 0.26% to 0.32% on Fair Isle and from 0.31% to 0.43% when all non-hybrid individuals are considered. The total nucleotide diversity over all individuals (and species, H_t_) is 0.53%. Differences between heterozygosity and population nucleotide diversity of one order of magnitude are observed even on Fair Isle (top panel of Figure 6A), where the aggregation of populations characterised by geographic isolation by distance is unlikely to play a role. Rather, these differences are likely to be the result of high levels of selfing. The per-nucleotide net divergence between species (π_(net)_ or d_a_) ranges from 0.07% (*E. arctica* – *E. foulaensis)* to 0.32% (*E. arctica* – *E. micrantha)* on Fair Isle, while this is lower (0.07% to 0.24%) in the broader comparison (both comparisons exclude hybrid individuals). With the exception of comparisons including Fair Isle *E. micrantha,* these estimates of between-species divergence are lower than the diversity found within each species (Figure 6A and B).

**Figure 6.**
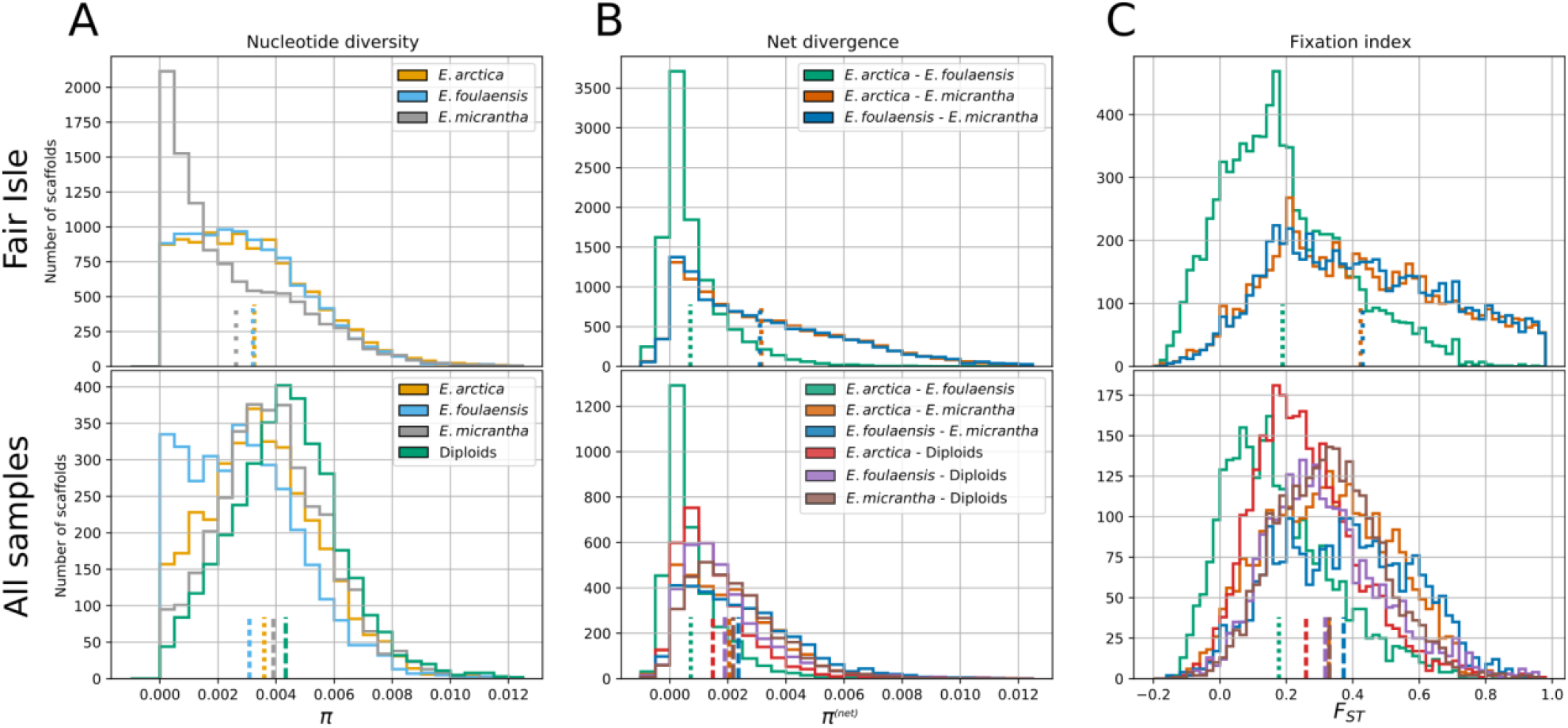
Considerable genetic structure with little differentiation between *Euphrasia* species. Histograms show per-scaffold statistics with population means indicated by dashed lines at the bottom of each panel. The top row shows results based on the tetraploid data set for non-hybrid individuals from Fair Isle, while the bottom row is based on the conserved set and it includes all non-hybrid individuals (with all diploids treated as one group). A) Nucleotide diversity on Fair Isle is slightly lower in *E. micrantha* than in *E. arctica* and *E. foulaensis;* and overall these values are considerably higher than per-individual heterozygosity estimates (Figure 3B); consistent with high levels of selfing. B) On Fair Isle, the net divergence shows very similar patterns in the comparisons involving *E. micrantha,* while divergence between *E. arctica* and *E. foulaensis* is much lower. With wider sampling the divergence between *E. micrantha* and the other species is lower, an indication that mainland *E. micrantha* carries alleles shared with the other species. The net divergences estimated here are of similar magnitude or lower than the nucleotide diversities shown in (A). C) While net divergence tends to be low between species of *Euphrasia,* the fixation index can be high, for example Fair Isle comparisons including *E. micrantha.* This genetic differentiation disappears when samples from additional populations are included, indicating that allelic frequency divergence is greater on population level than on the species level.

Our finding of clear genetic clusters but low divergence suggests that species may be recent, however these analyses do not indicate the amount of gene flow. We tested the extent of allele frequency differences, as measured by the fixation index (F_ST_), and found considerable genetic structure between Fair Isle *E. arctica* and *E. foulaensis* on one hand, and *E. micrantha* on the other. In comparisons to *E. micrantha,* genetic structure is high with mean F_ST_ values of 0.44 (*E. arctica-E. micrantha)* and 0.43 (*E. foulaensis-E. micrantha)*, while 11% of scaffolds show F_ST_>0.8 and 38% of scaffolds have F_ST_>0.5. Both F_ST_-distributions have very similar shapes (Figure 5C). Genetic structure is lower in the comparison between *E. arctica* and *E. foulaensis,* with a mean F_ST_ of 0.21, and only a minority of scaffolds with high values (0.11 %>0.8; 7.4%>0.5). On a larger scale, including all nonhybrid individuals and treating the diploids as one group, the differentiation between species is lower and is more similar across comparisons, with mean F_ST_ values ranging from 0.18 to 0.37, with the largest value for the comparison *E. foulaensis-E. micrantha* (Figure 6C, bottom panel). This is in line with a previous study that showed genetic variation in the related species *E. arctica* and *E. nemorosa* is structured by region of origin more than by species (French et al., 2008).

### Genetic relationships cannot be explained by a single tree

To characterise genome-wide relationships of populations and species, we generated trees from thousands of genomic scaffolds and used these to produce a consensus tree indicative of the relationships of populations and species. The ASTRAL consensus tree has a high posterior probability (PP, ≥0.97) for all nodes except for the clade of mainland polyploids (PP=0.82; Figure 7A). It does not generally show grouping by species, however, the diploids and Fair Isle *E. micrantha* are placed in distinct clades. The node ages, which are given in coalescence time, are comparatively young, the oldest being 1.39 (in generations scaled by 2N_e_, corresponding to a coalescence probability of 75%). Such recent divergence gives much opportunity for incomplete lineage sorting, i.e. the maintenance of ancestral polymorphism in present-day taxa. To better characterise the diverse genealogical histories present in the genome, we inspected the individual gene trees. Across 3454 scaffolds there was no clear congruence between individual trees and no general grouping by species, except some clustering of Fair Isle individuals of *E. micrantha* (M1-3; Figure 7B). This suggests that while broadscale relationships reflect divergence of Fair Isle *E. micrantha* from other tetraploids, and while diploids cluster separately, there is substantial gene tree incongruence and phylogenetic complexity.

**Figure 7.**
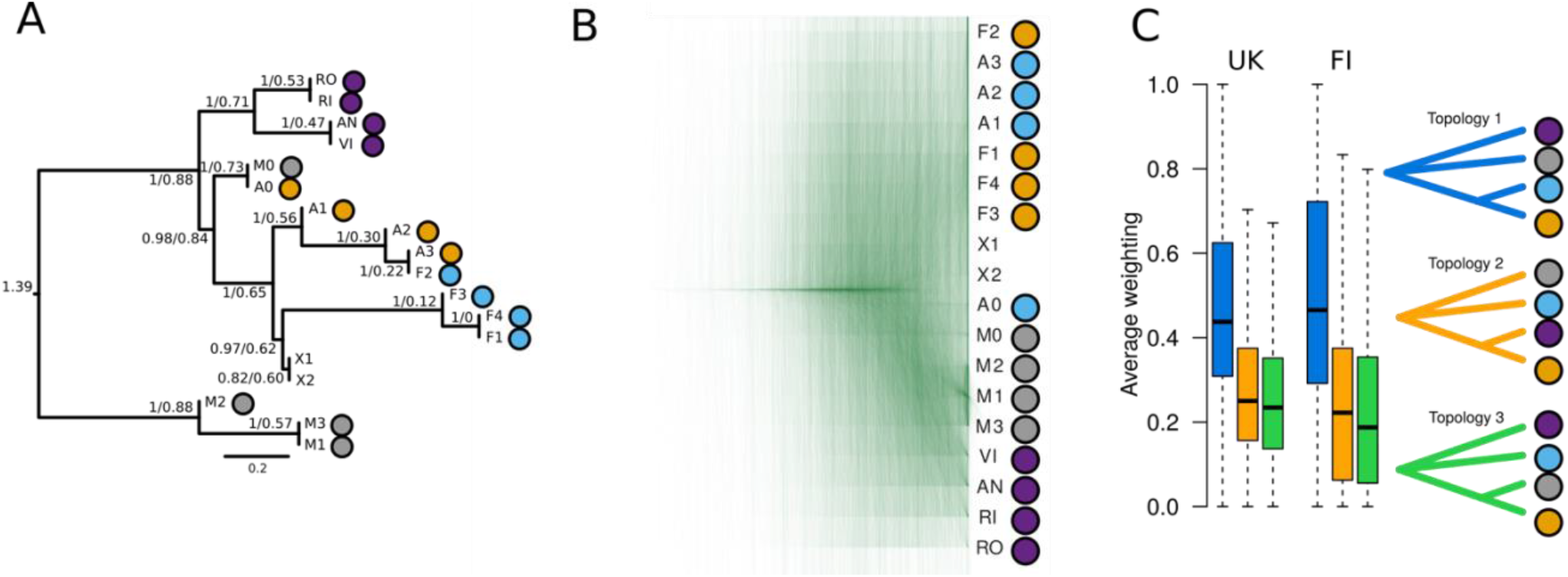
Complex evolutionary relationships and extensive discordance in *Euphrasia.* A) ASTRAL consensus tree based on 3454 per-scaffold trees from the conserved scaffold set. The numbers are a node’s posterior probability and its age in coalescent units. The tree is rooted at the longest branch (between Fair Isle *E. micrantha* and all other individuals). B) Overlaid gene trees of the conserved scaffold set show no single clear species relationship. C) Topological weighting of 3454 trees of all Fair Isle non-hybrids (FI) and all non-hybrid tetraploids (UK), both using the diploids as an outgroup, carried out with Twisst. While blue topology 1 tends to receive the highest weighting, few trees have a very high weight (near 1) for any one topology. The two alternative topologies receive similar levels of support. Coloured dots represent taxa, to match Figure 3.

To further categorise and quantify the different trees present in the genome, and to detect possible routes of gene flow, we conducted topology weighting with Twisst (Martin and van Belleghem, 2017). This compares individual trees to a set of reference topologies, where here we test the three possible unrooted topologies of four groups (each tetraploid species, excluding hybrids, and all diploids as an outgroup). Using the Fair Isle individuals, “Topology 1” (*E. arctica* and *E. foulaensis* as closest relatives, *E. micrantha* on a separate branch) received the highest mean weight of 51.23%. Both alternative topologies have similar mean weights of 23.40% and 25.37, see Figure 7C. Few of the 3454 per-scaffold trees matched one of the three reference topologies (419, 15, and 10, respectively). The topology mean weights were similar when mainland tetraploids were included (47.2%, 25.40%, and 27.48%, with 105, 3, and 1 trees matching the reference topologies entirely). For both, Fair Isle and the wider sampling, the topology with *E. arctica* and the diploids as closest relatives has a somewhat higher weight than the alternative (*E. micrantha* and *E. arctica* closest relatives), suggesting possible gene flow between *E. arctica* and the diploids, or between *E. micrantha* and *E. foulaensis.* We tested this with Patterson’s D statistic using the diploids as an outgroup and found that D only deviates significantly from zero (D=0.02, p=0.005) when the mainland tetraploids are included. This suggests gene flow between (mainland) *E. arctica* and the diploids, with an admixture fraction, measured as fG of 2.5% (Green et al., 2010). This value should be interpreted as additional admixture on top of any background level between all pairs of species.

## Discussion

Adaptive divergence is a major driver of speciation but remains poorly studied in polyploid organisms. In the present study, we use whole genome sequencing in combination with common garden experimental approaches to understand the nature of species differences in a complex tetraploid group. We have focused on an isolated island system where we can characterise genome-wide diversity and divergence of three differentially adapted eyebright species in sympatry. We discuss our findings regarding the role of divergent ecological selection in recent speciation, the history of subgenome evolution in polyploids, and the contribution of standing variation to adaptation in new lineages.

Studies of speciation genomics may gain valuable insights by investigating taxa at different stages of the speciation trajectory, from the earliest stages of divergence through to genetically differentiated species with strong reproductive barriers (Via, 2009; Twyford et al., 2014). We have shown that three species of eyebrights adapted to contrasting environments are characterised by a gradient of genomic differentiation, with grassland *E. arctica* and coastal *E. foulaensis* closely related, while the dry heathland specialist *E. micrantha* is genetically and morphologically more distinct. This parallels other genomic studies of adaptive divergence where species boundaries show different degrees of permeability, underpinned by different reproductive isolating barriers (Peccoud et al., 2009; Dasmahapatra et al., 2012; Renaut et al., 2012; Nosil et al., 2012; Roda et al., 2013). Reproductive isolation between *E. arctica* and *E. foulaensis* is incomplete and there is evidence of ongoing gene flow. However, the long tail in the pairwise F_ST_ distribution reveals many differentiated scaffolds, which may represent important regions of divergence involved in adaptation to harsh coastal environments (Lowry et al., 2008; Lyu et al., 2018; Dong et al., 2020), and the maintenance of species identities (Nosil et al., 2009; Ravinet et al., 2017). In contrast, *E. micrantha* shows genome-wide divergence from the other two taxa, despite a similar degree of geographic co-occurrence. Here, the high selfing rate of *E. micrantha* (Stone, 2013), and intrinsic postzygotic barriers (genomic incompatibilities), are likely to underlie reproductive isolation. However, our genomic analyses suggest the genetic distinctness of *E. micrantha* may be a feature of the Fair Isle population rather than the species as a whole. This could be a consequence regional differentiation caused by strong drift and the local fixation of genomic incompatibilities, local hybridisation, or polytopic origins as seen in other polyploids (Schwarzbach and Rieseberg, 2002; Soltis et al., 2004; Ainouche et al., 2012; Lowe and Abbott, 2015).

Key to understanding polyploid histories in British eyebrights, and many other postglacial groups such as octo- and dodecaploid *Cerastium* (Brysting et al., 2007), tetra-hexaploid species of *Silene* (Popp et al., 2005), and the highly variable ploidy *Primula* section *Aleuritia* (Guggisberg et al., 2006), is the use of polyploid-aware analyses. Our *k*-mer based polyploid tool Tetmer effectively estimates heterozygosity within and divergence between sub-genomes of polyploids directly from the reads, without the need for genome assembly. In *Euphrasia,* this approach revealed considerable between individual variation in heterozygosity within a species, indicative of a mixed mating system. This approach of analysing divergence within an individual proves complementary to comparative genomics between diploid and tetraploids, which revealed a set of scaffolds shared between diploids and tetraploids. This points to a (relative of) a British diploid acting as a parental progenitor to the British tetraploids, with the origin of the other, highly divergent, sub-genome (~5% divergence), unknown.

The absence of a second diploid ancestor and distinct genetic clustering of the diploids suggests that allopolyploidy in *Euphrasia* is not recent. This proves problematic for dating the polyploidy event(s), which would require extant but genetically isolated diploid relatives (Doyle and Egan, 2010). A further complication is that ongoing gene flow, evidenced by plastid genome sharing and the presence of natural diploid-tetraploid hybrids (Yeo, 1956), is blurring the split between the diploids and tetraploids. Polyploid species tend to go through a process of diploidisation, involving reduction in chromosome number, homeologous rearrangements, and gene losses leaving paleopolyploids with pre-polyploidy genome sizes (Mandáková and Lysak, 2018). This process does not seem to progress at equal pace in different polyploids. For instance, *Brassica napus,* an allopolyploid formed 7500-12,500 years ago and which has extant diploid relatives, experienced extensive homeologous rearrangements (Chalhoub et al., 2014), while teff *(Eragrostis tef,* VanBuren et al., 2020), formed 1.1 million years ago without known diploid ancestors, does not show signs of large-scale rearrangements. Whether homeologous exchange reshaped the sub-genomes of allotetraploid *Euphrasia,* and whether these potential rearrangements are shared between species and ploidy levels is likely to have major effects on interspecies gene flow, divergences, and ultimately on adaptation and speciation. These questions of sub-genome structure and genomic rearrangements will be addressed with an improved genome assembly, generated by the Darwin Tree of Life project (https://www.darwintreeoflife.org), and further population sequencing.

Our data provide insights into the enigmatic relationships of eyebrights, showing that their speciation history has been shaped by allopolyploidy with consistent sub-genome divergence, that overall species divergence is extremely shallow, and that no single phylogenetic tree can represent their evolutionary history. Since the wide adoption of genome-scale data for the study of adaptive radiations it has become clear that many groups are characterised by reticulate evolution, and analysing the reticulate evolution that gave rise to the diverse forms observed has become a main focus of speciation biology (Rieseberg et al., 2003; Cui et al., 2013; Malinsky et al., 2018; Sun et al., 2018; Feng et al., 2019). Because adaptation is underpinned by advantageous variants, adaptation in the absence of gene flow is expected to involve long waiting times for such variants to evolve. Like in many other rapid radiations, the relationships of tetraploid eyebrights are reticulate and reproductive isolation between them is incomplete, similar to tetraploid species of Arabidopsis (Jørgensen et al., 2011). In such settings it is easy for adaptive variants to cross species boundaries (Morjan and Rieseberg, 2004), generating novel phenotypic and genic combinations, which may lead to differential adaptation and reproductive isolation (Pardo-Diaz et al., 2012). Partial selfing may exacerbate this, as is seen in *Epipactis* orchids (Squirrell et al., 2002). This diversification process, termed “combinatorial speciation” (Marques et al., 2019), is very likely to underlie adaptation and the formation of taxonomic complexity in the young, and thus mutation-limited group of tetraploid *Euphrasia.*

## Methods

### Phenotypic differentiation

#### Natural populations

To establish the extent of phenotypic species differences, we measured morphological traits for *Euphrasia arctica, E. foulaensis* and *E*. *micrantha* in natural populations on Fair Isle. For each of two populations per species (Supplemental table 1) we measured 30 individual plants in the field at a single census point. The set of seven traits we measured is commonly used in *Euphrasia* identification (Metherell and Rumsey, 2018) and includes: plant height (Height), corolla length, number of nodes below the first flower (Nodes to flower), length ratio of the internode below the first flower and the leaf subtending the first flower (Leaf/internode ratio), number of leaf teeth (Leaf teeth), capsule width, and capsule length. All length measurements were made in mm, to the nearest 0.1mm using digital callipers.

##### Common garden experiment

We collected seeds from wild-collected open-pollinated plants, pooling seeds within a population. We then placed seeds in individual pots filled with a peat-free soil mix, which is a bark-based substrate of neutral pH (RGBE1), in December. The pots were kept outside at the Royal Botanic Garden Edinburgh (RGBE). We recorded germination daily from the beginning of April and supplied *Euphrasia* with host plants (twelve species, sourced from Fair Isle and commercial seed suppliers, see Supplemental table 3) at intervals of about two weeks. At the time of first flowering, we measured the same traits as in natural populations. At the end of September (or at the time of death), for each plant, we recorded its final height, the number of reproductive nodes (a measure of fitness), and length and width of one capsule. We also recorded which plants had died before the end of the experiment.

##### Morphological trait analysis and data visualisation

Trait differences were assessed with linear mixed effect models (R version 3.6.1 package lme4, https://github.com/lme4/lme4/). In the common garden study, host species was used as a random effect and transplant time was included as a covariate if the model’s log likelihood improved significantly. When analysing trait differences between species, genotype was included as a (nested) random effect. The significance of trait differences between populations or species was assessed with general linear hypothesis test (function ‘glht’) as implemented in the R package ‘multcomp’ (version 1.4-10). The p-values of trait differences were corrected for multiple tests on the level of each trait (rows in Fig. 2A).

To visualise individual clustering by phenotype, we carried out PCA with R’s built-in function ‘princomp’. To assess how well individuals can be classified into species, we ran linear discriminant analyses (LDAs), function ‘lda’ of the R package ‘MASS’, version 7.3-51.4). Classification was done with cross-validation (where each individual is, in turn, left out of the data matrix and classified according to its trait values compared to all other individuals of known species identity). For plotting, we ran ‘lda’ without cross-validation and used the function ‘predict’ to obtain two scores for each individual, which were plotted in two dimensions.

#### Genomic sequencing and analyses

##### Sample processing and sequencing

We collected individual *Euphrasia* plants in the field into desiccating silica gel. After grinding samples with a tissue mill using ceramic beads, we extracted DNA using Qiagen DNeasy Plant Mini Kit following the manufacturer’s instructions. DNA extraction for the *E. arctica* reference sample was performed by AmpliconExpress using their in-house high molecular weight DNA extraction protocol.

We generated short-read Illumina sequence data and long-read PacBio data for a range of *Euphrasia* samples. Full details of the sequencing protocols and specifications are provided in Supplemental Table 4.

##### Plastid genome and rDNA analyses

Plastid genomes were assembled from forward and reverse Illumina reads of each *Euphrasia* sample, using Novoplasty. Assembled plastids were manually curated and edited to give a standard order of the large single copy (LSC), inverted repeat 1 (IR1), small single copy (SSC) and IR2 using Geneious v11.1. Genome annotation of individual *E. arctica* A1 was performed using DOGMA (Wyman et al., 2004) with manual editing and curation, with annotation carried over to other samples using the ‘annotate from database’ option in Geneious. Plastid genomes were aligned using MAFFT. Phylogenetic analyses were performed using IQ-TREE (Nguyen et al., 2014), with the best evolutionary model inferred using model fitting and model assessment based on Bayesian Information Criterion, and level of branch support inferred via 1000 rapid bootstrap replicates. To help infer the direction of evolutionary change, we included an unpublished plastid assembly of the divergent *E. antarctica* (Twyford and Ness, unpublished) in the initial alignment and tree building, used this sample to root the phylogeny, and subsequently removed this long branch for better tree visualisation. The final alignment of British *Euphrasia* samples included 144,899 constant sites, 174 parsimony-informative and 133 singleton sites.

rDNA was assembled from the same read data as described above, but setting the expected assembly size to 9000-20,000bp and using a 1380 bp seed sequence of the rDNA cluster, obtained from a run of the RepeatExplorer pipeline (Novák et al., 2013). The assembler produced variable results, with some species having fully assembled circularised arrays, while some had multiple contigs. Initial comparisons of assembled arrays suggested some taxa were not alignable outside the rDNA coding region (results not shown). We therefore trimmed assemblies to the 5.8Kbp rDNA coding region (comprising 18S, ITS1, 5.8S, ITS2, 26S). Pairwise alignments were performed with MAFFT, and a neighbour joining tree constructed using Geneious.

##### K-mer analyses

We modelled the shape of *k*-mer spectra based on the infinite alleles model. Diploids and autotetraploids were treated like samples of two or four genomes from a single population. Allotetraploids were treated like two samples from each of two diverged populations. The corresponding formulae were derived using the block-wise site-frequency spectrum framework (Lohse et al., 2011; Lohse et al., 2016). Supplemental Text 2 describes this in more detail.

All models are implemented in our app, Tetmer, which can be used to fit parameters to *k*-mer spectra and which is available from GitHub (https://github.com/hannesbecher/shiny-k-mers).

##### Mapping and variant analysis

We generated a reference assembly of *E. arctica* from Illumina paired-end and PacBio data using the hybrid assembler SPAdes (Bankevich et al., 2012). The assembly was polished with ntEdit (Warren et al., 2019). The *k*-mer completeness of the assembly, as compared to the sequencing data, was assessed with the *k*-mer analysis toolkit (Mapleson et al., 2016). Inspection with blobtools (Laetsch and Blaxter, 2017) revealed that there were no contaminations at sequencing depths and CG-contents similar to *Euphrasia* DNA. We removed all scaffolds with an average mapping depth < 40x (inspection with blobtools had shown that this threshold separated contaminations from target scaffolds).

We mapped short-read data generated from natural populations to the reference using bwa-mem (Li 2013, https://arxiv.org/abs/1303.3997v2), and we computed per-scaffold mapping depths. These computations were automated using Snakemake pipelines (Köster and Rahmann, 2012). We then compared the mapping depth information across individuals. Two sets of scaffolds were selected for variant calling: the universal data set, which comprises 3454 scaffolds with disomic coverage in all individuals, and the tetraploid set, which comprises the universal set and 7,189 more scaffolds restricted to tetraploid individuals (totalling 10,643 scaffolds). All other scaffolds of the reference were concatenated and kept in the assembly to avoid non-specific mapping of reads from diverged regions to our scaffold set. We called variants with freebayes (Garrison and Marth, 2012, https://arxiv.org/abs/1207.3907) separately for the universal and tetraploid scaffold sets. In the resulting VCF files, we used only biallelic SNPs with a quality value greater than 100. The VCF files where handled, interactively, in jupyter lab (https://github.com/jupyterlab) using the scikit-allel package (https://doi.org/10.5281/zenodo.3238280). An HTML report of the analyses is included in the zenodo archive published alongside this article (https://doi.org/10.5281/zenodo.3774489).

##### Analysis of population structure

We used two complementary approaches to infer population structure. To avoid spurious signal from sites in linkage disequilibrium in these analyses, we selected one random polymorphism per scaffold from the “conserved” data set. We then carried out PCA as implemented in the adegenet package, version 2.1.2 (Jombart and Ahmed, 2011) and we ran FastStructure (Raj et al., 2014) with *K* values ranging from 1 to 10. As FastStructure may underestimate the extent of admixture, we then analysed the same dataset in STRUCTURE (Pritchard et al., 2000), using a subset of *K* values (k=1-5) that were deemed to be most likely. STRUCTURE was run with the admixture model, using 3 replicates per k-value, with 100,000 burn-in generations followed by 100,000 MCMC generations. The optimal number of clusters was inferred using the ad hoc statistic delta *K* (Evanno et al., 2005). We combined multiple STRUCTURE runs in CLUMPP (Jakobsson and Rosenberg, 2007) and visualised STRUCTURE plots with a custom approach.

##### Tree-based analyses

We generated per-scaffold unrooted neighbour-joining trees based on pairwise estimates of net-nucleotide divergence between individuals using the nj function of the R package ape, version 5.3 (Paradis and Schliep, 2018). These trees were used for a number of downstream analyses, as follows. We generated a consensus tree using ASTRAL (Zhang et al., 2018). We rooted the tree at the longest branch, which connected Fair Isle *E. micrantha* to all the other samples. We then performed topology weighting as implemented in the package Twisst (Martin and van Belleghem, 2017). As candidate topologies, we used all three possible trees with four groups (diploid outgroup and one group per species of *Euphrasia).* We run Twisst twice, once on the Fair isle samples and once including the mainland tetraploids. Finally, we carried out two sets of ABBA-BABA tests with Dsuite (Malinsky et al., https://doi.org/10.1101/634477), using all diploids as the outgroup. First, we used all non-hybrid individuals from Fair Isle, grouped by species, and second, we added the mainland tetraploids according to their species.

## Supporting information

Concatenated supplemental items

## Author contributions

H.B. and A.T. designed the experiments. H.B., A.T., G.P., and M.B. set up the experiments. All authors collected the data. C.M. confirmed the species identifications. N.R. provided valuable support on Fair Isle. H.B. analysed the data. H.B and A.T. wrote the manuscript. All authors commented on and approved the manuscript.

## Acknowledgements

This work was funded by NERC grants (NE/R010609/1; NE/L011336/1; NE/N006739/1) awarded to A.T. We are extremely grateful to the Fair Isle Bird Observatory for providing accommodation and infrastructure, and the RBGE for providing plant growing facilities. We thank Edinburgh Genomics for generating Illumina data and the Centre for Genomic Research at the University of Liverpool for generating PacBio data. We also wish to thank Cristina Rosique for her help setting up the common garden experiment, and Mabon Elis for help with Figure 1. We thank Richard Ennos and Simon Martin who commented on an earlier version of the manuscript. Kamil Jaron supplied the smudge plots of the four diploids, for which we thank hi.

