## Supplementary material for "Maintenance of species differences in closely related tetraploid parasitic *Euphrasia* (Orobanchaceae) on an isolated island": Concatenated supplemental items

#### SUPPLEMENTAL FIGURE 1

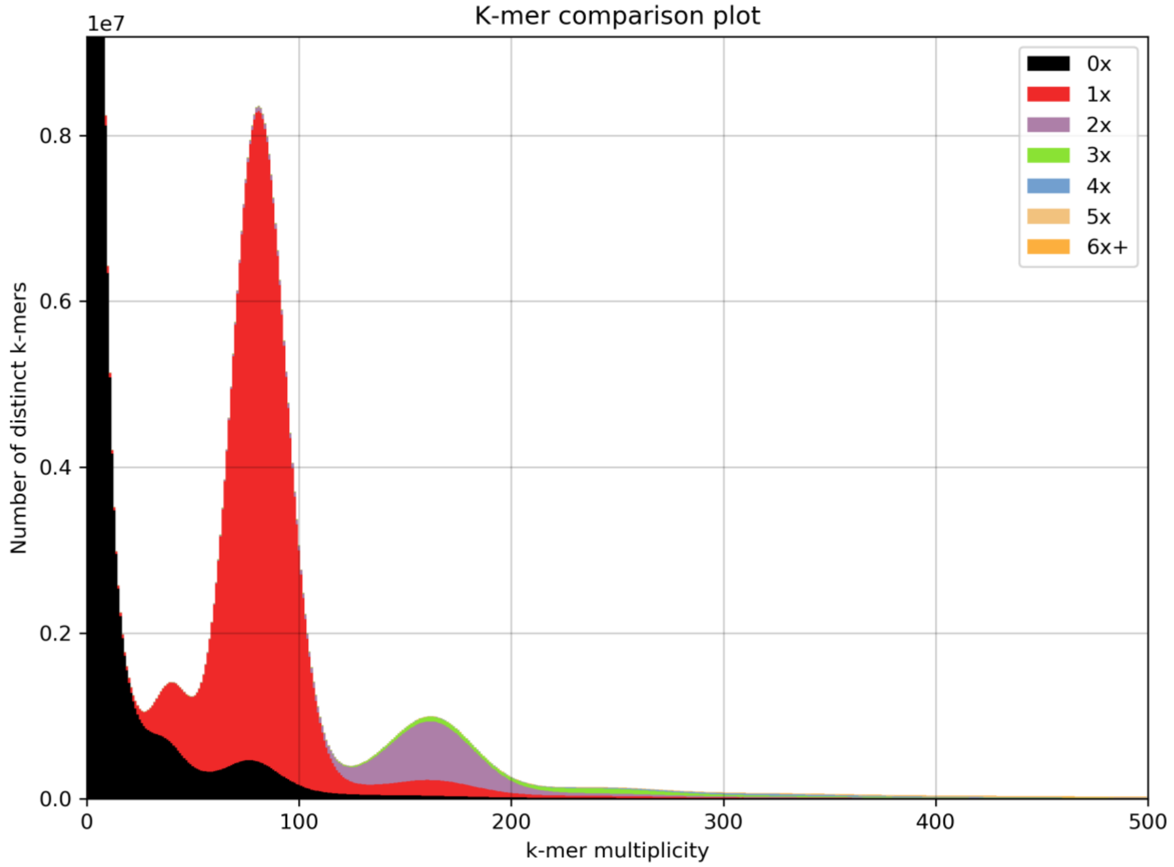

A k-mer completeness plot for our assembly of *E. arctica* generated with KAT.

#### SUPPLEMENTAL FIGURE 2

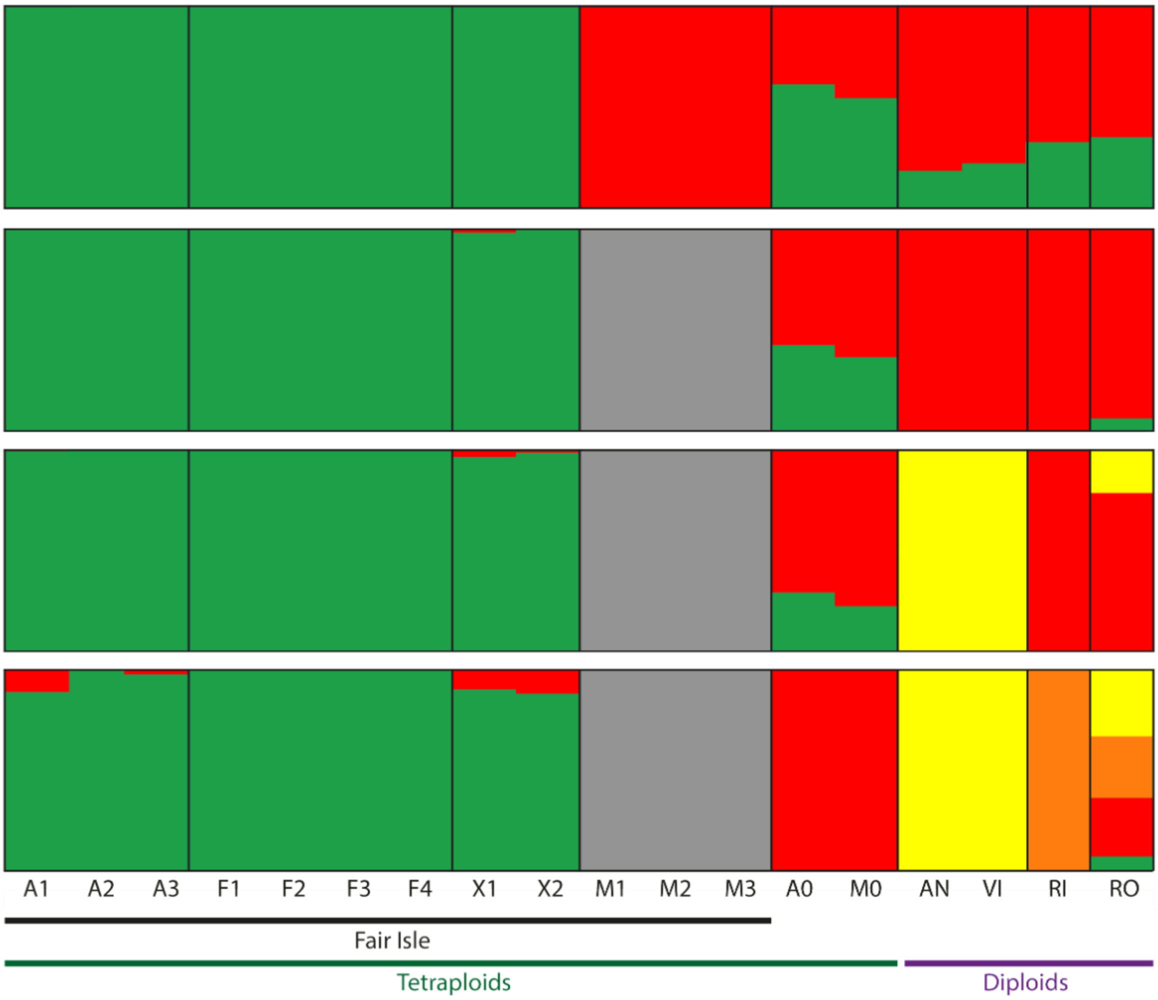

Output of STRUCTURE runs with K values 2-5.

#### SUPPLEMENTAL TABLE 1

Locations and descriptions of populations sampled on Fair Isle.

| | Population code | latitude | longitutde | Description |
|---|---|---|---|---|
| E. arctica | G1 | 59.524622 | -1.636012 | Road side, near school |
| E. arctica | G2 | 59.517135 | -1.64082 | Road side, near chapel |
| E. foulaensis | C1 | 59.535645 | -1.602904 | Coastal turf, Buness peninsula |
| E. foulaensis | C2 | 59.513855 | -1.650254 | Coastal turf, South light |
| E. micrantha | H1 | 59.537202 | -1.629517 | Heath land, North of airstrip |
| E. micrantha | H2 | 59.548057 | -1.615796 | Heath land, by bend in road to N light |

#### SUPPLEMENTAL TABLE 2

Means and standard errors of trait measurements from natural populations (Fair Isle) and from the common garden.

Submitted separately.

#### SUPPLEMENTAL TABLE 3

Origins of host seed used in the common garden experiment.

| Species | Source | Notes |
|---|---|---|
| *Holcus lanatus* | Emorsgate Seeds* | Seeds ordered in early 2019 |
| *Lolium perenne* | Emorsgate Seeds* | Seeds ordered in early 2019 |
| *Trifolium repens* | Emorsgate Seeds* | Seeds ordered in early 2019 |
| *Armeria maritima* | Scotia Seeds** | Seeds ordered in early 2019 |
| *Juniperus vulgaris* | Fair Isle, wild collected | cuttings taken in September 2018 |
| *Calluna vulgaris* | Fair Isle, wild collected | cuttings taken in September 2018 |
| *Plantago maritima* | Fair Isle, wild collected | seeds collected in August 2018 |
| *Plantago lanceolata* | Fair Isle, wild collected | seeds collected in August 2018 |
| *Plantago coronopus* | Fair Isle, wild collected | seeds collected in August 2018 |
| *Rumex crispus* | Fair Isle, wild collected | seeds collected in August 2018 |
| *Luzula multiflora* | Fair Isle, wild collected | seeds collected in August 2018 |
| *Rumex acetosa* | Fair Isle, wild collected | seeds collected in August 2018 |
| * EMORSGATE SEEDS, Limes Farm, Tilney All Saints, King's Lynn, Norfolk, PE34 4RT | ** Scotia Seeds, Mavisbank, Farnell, Brechin, Angus, DD9 6TR | |

#### SUPPLEMENTAL TABLE 4

Individuals sequenced, heterozygosity and sub-genome divergence estimates.

Submitted separately.

### Supplemental text 1 – Magnitude and significance of trait value differences

#### Order of traits

The order of traits in the following tables corresponds to Figure 1D. Traits marked with * were recorded from natural populations.

1. Time transplant to flower in days
2. Time of 1st flower in Julian days
3. Height at 1st flower in mm
4. *Final height in mm
5. Number of reproductive nodes
6. *Corolla length in mm
7. *Number of nodes blow the flower
8. *Ration of the length of the leaf subtending the lowest flower and the internode beneath
9. *Number of leaf teeth
10. *Capsule width in mm
11. *Capsule length in mm
12. Germination rate
13. Proportion flowering of those germinated
14. Proportion early death out of those germinated

#### Trait differences in natural populations, species level

```
4
        Simultaneous Tests for General Linear Hypotheses

Fit: lmer(formula = log(height) ~ Category + (1 | Population), data = datcomp)

Linear Hypotheses:
                       Estimate Std. Error t value Pr(>|t|)
arc - fou == 0  1.39687    0.08663  16.125  0.00127 **
arc - mic == 0  0.71272    0.08663   8.227  0.00779 **
fou - mic == 0 -0.68415    0.08663  -7.897  0.00846 **
---
Signif. codes:  0 '*' 0.001 '' 0.01 '*' 0.05 '.' 0.1 ' ' 1
(Adjusted p values reported -- single-step method)
```

```
6

        Simultaneous Tests for General Linear Hypotheses

Fit: lmer(formula = corolla.size ~ Category + (1 | Population), data = datcomp)

Linear Hypotheses:
                       Estimate Std. Error t value Pr(>|t|)
arc - fou == 0   1.7196     0.1312  13.105  0.00193 **
arc - mic == 0   2.9225     0.1483  19.706  < 0.001 ***
fou - mic == 0   1.2029     0.1495   8.045  0.00807 **
---
Signif. codes:  0 '*' 0.001 '' 0.01 '*' 0.05 '.' 0.1 ' ' 1
(Adjusted p values reported -- single-step method)
```

7

```
        Simultaneous Tests for General Linear Hypotheses

Fit: lmer(formula = nd.to.fl ~ Category + (1 | Population), data =
datcomp[!rownames(datcomp) %in%
       c("17", "113"), ])

Linear Hypotheses:
                      Estimate Std. Error t value Pr(>|t|)
arc - fou == 0   1.5556     0.5485   2.836   0.1280
arc - mic == 0   2.4028     0.5491   4.376   0.0442 *
fou - mic == 0   0.8472     0.5480   1.546   0.3897
---
Signif. codes:  0 '*' 0.001 '' 0.01 '*' 0.05 '.' 0.1 ' ' 1
(Adjusted p values reported -- single-step method)
```

8

```
        Simultaneous Tests for General Linear Hypotheses

Fit: lmer(formula = log(lir) ~ Category + (1 | Population), data = datcomp)

Linear Hypotheses:
                      Estimate Std. Error t value Pr(>|t|)
arc - fou == 0  -0.7981     0.2801  -2.850   0.1266
arc - mic == 0   0.7305     0.2803   2.606   0.1540
fou - mic == 0   1.5286     0.2798   5.464   0.0244 *
---
Signif. codes:  0 '*' 0.001 '' 0.01 '*' 0.05 '.' 0.1 ' ' 1
(Adjusted p values reported -- single-step method)
```

9

```
        Simultaneous Tests for General Linear Hypotheses

Fit: lmer(formula = leaf.teeth ~ Category + (1 | Population), data = datcomp)

Linear Hypotheses:
                      Estimate Std. Error t value Pr(>|t|)
arc - fou == 0   1.1742     0.5699   2.060    0.245
arc - mic == 0   1.5244     0.5702   2.674    0.146
fou - mic == 0   0.3502     0.5697   0.615    0.823
(Adjusted p values reported -- single-step method)
```

10

```
        Simultaneous Tests for General Linear Hypotheses

Fit: lmer(formula = capsule.width ~ Category + (1 | Population), data =
datcomp[!rownames(datcomp) %in%
       c("17", "113"), ])

Linear Hypotheses:
                      Estimate Std. Error t value Pr(>|t|)
arc - fou == 0  -0.1279     0.3423  -0.374    0.928
arc - mic == 0   0.5693     0.3410   1.669    0.348
fou - mic == 0   0.6972     0.3419   2.039    0.250
(Adjusted p values reported -- single-step method)
```

11

```
        Simultaneous Tests for General Linear Hypotheses
```

```
Fit: lmer(formula = capsule.width ~ Category + (1 | Population), data =
datcomp[!rownames(datcomp) %in%
       c("17", "113"), ])

Linear Hypotheses:
                       Estimate Std. Error t value Pr(>|t|)
arc - fou == 0  -0.1279     0.3423  -0.374    0.928
arc - mic == 0   0.5693     0.3410   1.669    0.348
fou - mic == 0   0.6972     0.3419   2.039    0.250
(Adjusted p values reported -- single-step method)
```

#### TRAIT DIFFERENCES IN NATURAL POPULATIONS, POPULATION LEVEL

4

```
        Simultaneous Tests for General Linear Hypotheses

Fit: lm(formula = log(height) ~ Population, data = datcomp)

Linear Hypotheses:
                     Estimate Std. Error t value Pr(>|t|)
G1 - G2 == 0 -0.04677    0.07041  -0.664   0.9856
G1 - C2 == 0  1.41311    0.07041  20.070   <0.001 ***
G1 - C1 == 0  1.33386    0.07041  18.944   <0.001 ***
G1 - H2 == 0  0.78494    0.07041  11.148   <0.001 ***
G1 - H1 == 0  0.59373    0.07041   8.432   <0.001 ***
G2 - C2 == 0  1.45988    0.07041  20.734   <0.001 ***
G2 - C1 == 0  1.38063    0.07041  19.608   <0.001 ***
G2 - H2 == 0  0.83171    0.07041  11.812   <0.001 ***
G2 - H1 == 0  0.64050    0.07041   9.097   <0.001 ***
C2 - C1 == 0 -0.07925    0.07041  -1.126   0.8702
C2 - H2 == 0 -0.62817    0.07041  -8.922   <0.001 ***
C2 - H1 == 0 -0.81938    0.07041 -11.637   <0.001 ***
C1 - H2 == 0 -0.54892    0.07041  -7.796   <0.001 ***
C1 - H1 == 0 -0.74013    0.07041 -10.512   <0.001 ***
H2 - H1 == 0 -0.19121    0.07041  -2.716   0.0772 .
---
Signif. codes:  0 '*' 0.001 '' 0.01 '*' 0.05 '.' 0.1 ' ' 1
(Adjusted p values reported -- single-step method)
```

6

```
        Simultaneous Tests for General Linear Hypotheses

Fit: lm(formula = corolla.size ~ Population, data = datcomp)

Linear Hypotheses:
                     Estimate Std. Error t value Pr(>|t|)
G1 - G2 == 0  0.08214    0.15825   0.519    0.995
G1 - C2 == 0  1.61671    0.16293   9.923   <0.001 ***
G1 - C1 == 0  1.89286    0.15825  11.961   <0.001 ***
G1 - H2 == 0  3.06071    0.20430  14.981   <0.001 ***
G1 - H1 == 0  2.89756    0.17600  16.464   <0.001 ***
G2 - C2 == 0  1.53457    0.16293   9.419   <0.001 ***
G2 - C1 == 0  1.81071    0.15825  11.442   <0.001 ***
G2 - H2 == 0  2.97857    0.20430  14.579   <0.001 ***
G2 - H1 == 0  2.81541    0.17600  15.997   <0.001 ***
```

```
C2 - C1 == 0  0.27614    0.16293   1.695    0.533
C2 - H2 == 0  1.44400    0.20794   6.944   <0.001 ***
C2 - H1 == 0  1.28084    0.18021   7.107   <0.001 ***
C1 - H2 == 0  1.16786    0.20430   5.716   <0.001 ***
C1 - H1 == 0  1.00470    0.17600   5.709   <0.001 ***
H2 - H1 == 0 -0.16316    0.21833  -0.747    0.975
---
Signif. codes:  0 ‘*’ 0.001 ‘’ 0.01 ‘*’ 0.05 ‘.’ 0.1 ‘ ’ 1
(Adjusted p values reported -- single-step method)
```

7

```
        Simultaneous Tests for General Linear Hypotheses

Fit: lm(formula = nd.to.fl ~ Population, data = datcomp[!rownames(datcomp) %in%
       c("17", "113"), ])

Linear Hypotheses:
                     Estimate Std. Error t value Pr(>|t|)
G1 - G2 == 0   1.2881     0.3719   3.463  0.00864 **
G1 - C2 == 0   2.3548     0.3719   6.331  < 0.001 ***
G1 - C1 == 0   2.0548     0.3719   5.525  < 0.001 ***
G1 - H2 == 0   2.9214     0.3719   7.855  < 0.001 ***
G1 - H1 == 0   3.1835     0.3750   8.489  < 0.001 ***
G2 - C2 == 0   1.0667     0.3654   2.919  0.04525 *
G2 - C1 == 0   0.7667     0.3654   2.098  0.29328
G2 - H2 == 0   1.6333     0.3654   4.469  < 0.001 ***
G2 - H1 == 0   1.8954     0.3686   5.142  < 0.001 ***
C2 - C1 == 0  -0.3000     0.3654  -0.821  0.96334
C2 - H2 == 0   0.5667     0.3654   1.551  0.63200
C2 - H1 == 0   0.8287     0.3686   2.248  0.22144
C1 - H2 == 0   0.8667     0.3654   2.372  0.17224
C1 - H1 == 0   1.1287     0.3686   3.062  0.03012 *
H2 - H1 == 0   0.2621     0.3686   0.711  0.98041
---
Signif. codes:  0 ‘*’ 0.001 ‘’ 0.01 ‘*’ 0.05 ‘.’ 0.1 ‘ ’ 1
(Adjusted p values reported -- single-step method)
```

8

```
        Simultaneous Tests for General Linear Hypotheses

Fit: lm(formula = log(lir) ~ Population, data = datcomp)

Linear Hypotheses:
                     Estimate Std. Error t value Pr(>|t|)
G1 - G2 == 0   0.3861     0.1105   3.493  0.00797 **
G1 - C2 == 0  -0.8379     0.1068  -7.846  < 0.001 ***
G1 - C1 == 0  -0.3711     0.1085  -3.419  0.01012 *
G1 - H2 == 0   0.7628     0.1095   6.965  < 0.001 ***
G1 - H1 == 0   1.0839     0.1085   9.985  < 0.001 ***
G2 - C2 == 0  -1.2240     0.1068 -11.462  < 0.001 ***
G2 - C1 == 0  -0.7572     0.1085  -6.976  < 0.001 ***
G2 - H2 == 0   0.3767     0.1095   3.440  0.00949 **
G2 - H1 == 0   0.6978     0.1085   6.428  < 0.001 ***
C2 - C1 == 0   0.4668     0.1047   4.457  < 0.001 ***
C2 - H2 == 0   1.6007     0.1057  15.140  < 0.001 ***
C2 - H1 == 0   1.9218     0.1047  18.350  < 0.001 ***
C1 - H2 == 0   1.1339     0.1075  10.548  < 0.001 ***
C1 - H1 == 0   1.4550     0.1065  13.660  < 0.001 ***
H2 - H1 == 0   0.3211     0.1075   2.987  0.03774 *
---
Signif. codes:  0 ‘*’ 0.001 ‘’ 0.01 ‘*’ 0.05 ‘.’ 0.1 ‘ ’ 1
(Adjusted p values reported -- single-step method)
```

9

```
         Simultaneous Tests for General Linear Hypotheses

Fit: lm(formula = leaf.teeth ~ Population, data = datcomp)

Linear Hypotheses:
                 Estimate Std. Error t value Pr(>|t|)
G1 - G2 == 0  0.85755    0.15126   5.669   <0.001 ***
G1 - C2 == 0  1.05385    0.14750   7.145   <0.001 ***
G1 - C1 == 0  2.15385    0.14993  14.366   <0.001 ***
G1 - H2 == 0  1.93162    0.15126  12.770   <0.001 ***
G1 - H1 == 0  1.97527    0.14993  13.175   <0.001 ***
G2 - C2 == 0  0.19630    0.14603   1.344    0.760
G2 - C1 == 0  1.29630    0.14848   8.730   <0.001 ***
G2 - H2 == 0  1.07407    0.14983   7.169   <0.001 ***
G2 - H1 == 0  1.11772    0.14848   7.528   <0.001 ***
C2 - C1 == 0  1.10000    0.14465   7.604   <0.001 ***
C2 - H2 == 0  0.87778    0.14603   6.011   <0.001 ***
C2 - H1 == 0  0.92143    0.14465   6.370   <0.001 ***
C1 - H2 == 0 -0.22222    0.14848  -1.497    0.667
C1 - H1 == 0 -0.17857    0.14713  -1.214    0.830
H2 - H1 == 0  0.04365    0.14848   0.294    1.000
---
Signif. codes:  0 '*' 0.001 '' 0.01 '*' 0.05 '.' 0.1 ' ' 1
(Adjusted p values reported -- single-step method)
```

10

```
         Simultaneous Tests for General Linear Hypotheses

Fit: lm(formula = capsule.width ~ Population, data = datcomp[!rownames(datcomp)
%in%
      c("17", "113"), ])

Linear Hypotheses:
                  Estimate Std. Error t value Pr(>|t|)
G1 - G2 == 0 -0.706981   0.094030  -7.519   <0.001 ***
G1 - C2 == 0 -0.537500   0.095276  -5.642   <0.001 ***
G1 - C1 == 0 -0.421627   0.099709  -4.229   <0.001 ***
G1 - H2 == 0  0.003929   0.086725   0.045   1.0000
G1 - H1 == 0  0.432005   0.089888   4.806   <0.001 ***
G2 - C2 == 0  0.169481   0.100690   1.683   0.5438
G2 - C1 == 0  0.285354   0.104895   2.720   0.0771 .
G2 - H2 == 0  0.710909   0.092641   7.674   <0.001 ***
G2 - H1 == 0  1.138986   0.095608  11.913   <0.001 ***
C2 - C1 == 0  0.115873   0.106013   1.093   0.8828
C2 - H2 == 0  0.541429   0.093905   5.766   <0.001 ***
C2 - H1 == 0  0.969505   0.096833  10.012   <0.001 ***
C1 - H2 == 0  0.425556   0.098400   4.325   <0.001 ***
C1 - H1 == 0  0.853632   0.101199   8.435   <0.001 ***
H2 - H1 == 0  0.428077   0.088434   4.841   <0.001 ***
---
Signif. codes:  0 '*' 0.001 '' 0.01 '*' 0.05 '.' 0.1 ' ' 1
(Adjusted p values reported -- single-step method)
```

11

```
         Simultaneous Tests for General Linear Hypotheses

Fit: lm(formula = capsule.width ~ Population, data = datcomp[!rownames(datcomp)
%in%
      c("17", "113"), ])

Linear Hypotheses:
                  Estimate Std. Error t value Pr(>|t|)
```

```
G1 - G2 == 0 -0.706981   0.094030  -7.519   <0.001 ***
G1 - C2 == 0 -0.537500   0.095276  -5.642   <0.001 ***
G1 - C1 == 0 -0.421627   0.099709  -4.229   <0.001 ***
G1 - H2 == 0  0.003929   0.086725   0.045   1.0000
G1 - H1 == 0  0.432005   0.089888   4.806   <0.001 ***
G2 - C2 == 0  0.169481   0.100690   1.683   0.5438
G2 - C1 == 0  0.285354   0.104895   2.720   0.0771 .
G2 - H2 == 0  0.710909   0.092641   7.674   <0.001 ***
G2 - H1 == 0  1.138986   0.095608  11.913   <0.001 ***
C2 - C1 == 0  0.115873   0.106013   1.093   0.8829
C2 - H2 == 0  0.541429   0.093905   5.766   <0.001 ***
C2 - H1 == 0  0.969505   0.096833  10.012   <0.001 ***
C1 - H2 == 0  0.425556   0.098400   4.325   <0.001 ***
C1 - H1 == 0  0.853632   0.101199   8.435   <0.001 ***
H2 - H1 == 0  0.428077   0.088434   4.841   <0.001 ***
---
Signif. codes:  0 ‘*’ 0.001 ‘’ 0.01 ‘*’ 0.05 ‘.’ 0.1 ‘ ’ 1
(Adjusted p values reported -- single-step method)
```

#### TRAIT DIFFERENCES IN THE COMMON GARDEN, SPECIES LEVEL

1

```
        Simultaneous Tests for General Linear Hypotheses

Fit: lmer(formula = Fd ~ Euphrasia.Taxon + (1 | gerGT) + (1 | Host),
       data = FdAll, REML = T)

Linear Hypotheses:
                       Estimate Std. Error t value Pr(>|t|)
arc - fou == 0    1.578      5.537   0.285    0.957
arc - mic == 0   -4.969      5.684  -0.874    0.690
fou - mic == 0   -6.547      5.716  -1.145    0.555
(Adjusted p values reported -- single-step method)
```

2

```
        Simultaneous Tests for General Linear Hypotheses

Fit: lmer(formula = Flower ~ Euphrasia.Taxon + (1 | gerGT) + (1 |
       Host), data = FdAll, REML = T)

Linear Hypotheses:
                       Estimate Std. Error t value Pr(>|t|)
arc - fou == 0  -0.8433     3.5089  -0.240    0.969
arc - mic == 0  -2.5972     3.6740  -0.707    0.777
fou - mic == 0  -1.7539     3.7106  -0.473    0.888
(Adjusted p values reported -- single-step method)
```

3

```
        Simultaneous Tests for General Linear Hypotheses

Fit: lmer(formula = log(Plant.height) ~ Euphrasia.Taxon + Transplant +
       (1 | gerGT) + (1 | Host), data = FdAll, REML = T)

Linear Hypotheses:
                       Estimate Std. Error t value Pr(>|t|)
arc - fou == 0   0.1487     0.1744   0.853    0.701
arc - mic == 0   0.2550     0.1782   1.431    0.432
fou - mic == 0   0.1062     0.1790   0.593    0.833
(Adjusted p values reported -- single-step method)
```

4

```
        Simultaneous Tests for General Linear Hypotheses

Fit: lmer(formula = log(Height) ~ Euphrasia.Taxon + Transplant + (1 |
       gerGT) + (1 | Host), data = FdAll, REML = T)

Linear Hypotheses:
                       Estimate Std. Error t value Pr(>|t|)
arc - fou == 0  0.25164    0.19112   1.317    0.478
arc - mic == 0  0.34798    0.19790   1.758    0.321
fou - mic == 0  0.09633    0.19926   0.483    0.884
(Adjusted p values reported -- single-step method)
```

5

```
        Simultaneous Tests for General Linear Hypotheses

Fit: lmer(formula = log(1 + Reproductive.nodes) ~ Euphrasia.Taxon +
       Transplant + (1 | gerGT) + (1 | Host), data = FdAll, REML = T)

Linear Hypotheses:
                       Estimate Std. Error t value Pr(>|t|)
arc - fou == 0 -0.09066    0.15157  -0.598    0.831
arc - mic == 0  0.41922    0.16520   2.538    0.163
fou - mic == 0  0.50987    0.16741   3.046    0.109
(Adjusted p values reported -- single-step method)
```

6

```
        Simultaneous Tests for General Linear Hypotheses

Fit: lmer(formula = Corolla.length ~ Euphrasia.Taxon + (1 | gerGT) +
       (1 | Host), data = FdAll, REML = T)

Linear Hypotheses:
                       Estimate Std. Error t value Pr(>|t|)
arc - fou == 0   0.4772     0.3837   1.244   0.5100
arc - mic == 0   1.7766     0.3879   4.580   0.0393 *
fou - mic == 0   1.2994     0.3888   3.342   0.0874 .
---
Signif. codes:  0 ‘*’ 0.001 ‘’ 0.01 ‘*’ 0.05 ‘.’ 0.1 ‘ ’ 1
(Adjusted p values reported -- single-step method)
```

7

```
        Simultaneous Tests for General Linear Hypotheses

Fit: lmer(formula = Nodes.to.flower ~ Euphrasia.Taxon + (1 | gerGT) +
       (1 | Host), data = FdAll, REML = T)

Linear Hypotheses:
                       Estimate Std. Error t value Pr(>|t|)
arc - fou == 0  -0.5544     0.6536  -0.848    0.704
arc - mic == 0   0.5287     0.6601   0.801    0.728
fou - mic == 0   1.0831     0.6617   1.637    0.359
(Adjusted p values reported -- single-step method)
```

8

```
        Simultaneous Tests for General Linear Hypotheses

Fit: lmer(formula = log(lir) ~ Euphrasia.Taxon + (1 | gerGT) + (1 |
```

```
       Host), data = FdAll, REML = T)

Linear Hypotheses:
                      Estimate Std. Error t value Pr(>|t|)
arc - fou == 0  -0.3040     0.3557  -0.855    0.700
arc - mic == 0   0.4369     0.3586   1.218    0.521
fou - mic == 0   0.7410     0.3592   2.063    0.245
(Adjusted p values reported -- single-step method)
```

9

```
        Simultaneous Tests for General Linear Hypotheses

Fit: lmer(formula = log(Number.of.leaf.teeth..exc..tip.) ~ Euphrasia.Taxon +
       Transplant + (1 | gerGT) + (1 | Host), data = FdAll)

Linear Hypotheses:
                      Estimate Std. Error t value Pr(>|t|)
arc - fou == 0 -0.02808    0.23120  -0.121    0.992
arc - mic == 0  0.12424    0.23278   0.534    0.861
fou - mic == 0  0.15232    0.23312   0.653    0.804
(Adjusted p values reported -- single-step method)
```

10

```
        Simultaneous Tests for General Linear Hypotheses

Fit: lmer(formula = Capsule.width ~ Euphrasia.Taxon + (1 | gerGT) +
       (1 | Host), data = FdAll, REML = T)

Linear Hypotheses:
                      Estimate Std. Error t value Pr(>|t|)
arc - fou == 0   0.1145     0.1379   0.831   0.7127
arc - mic == 0   0.7230     0.1474   4.904   0.0328 *
fou - mic == 0   0.6084     0.1479   4.113   0.0521 .
---
Signif. codes:  0 ‘*’ 0.001 ‘’ 0.01 ‘*’ 0.05 ‘.’ 0.1 ‘ ’ 1
(Adjusted p values reported -- single-step method)
```

11

```
        Simultaneous Tests for General Linear Hypotheses

Fit: lmer(formula = Capsule.Height ~ Euphrasia.Taxon + (1 | gerGT) +
       (1 | Host), data = FdAll, REML = T)

Linear Hypotheses:
                      Estimate Std. Error t value Pr(>|t|)
arc - fou == 0   0.7293     0.1311   5.562  0.00601 **
arc - mic == 0   1.6583     0.1766   9.388  < 0.001 ***
fou - mic == 0   0.9289     0.1789   5.193  0.00816 **
---
Signif. codes:  0 ‘*’ 0.001 ‘’ 0.01 ‘*’ 0.05 ‘.’ 0.1 ‘ ’ 1
(Adjusted p values reported -- single-step method)
```

12

```
        Simultaneous Tests for General Linear Hypotheses

Fit: glmer(formula = cbind(X1, X2) ~ spec + (1 | GT), data = ger,
       family = binomial)

Linear Hypotheses:
                      Estimate Std. Error z value Pr(>|z|)
```

```
arc - fou == 0   0.6520     0.6027   1.082    0.525  
arc - mic == 0   1.5402     0.6041   2.550    0.029 *
fou - mic == 0   0.8882     0.6039   1.471    0.305  
---
Signif. codes:  0 ‘*’ 0.001 ‘’ 0.01 ‘*’ 0.05 ‘.’ 0.1 ‘ ’ 1
(Adjusted p values reported -- single-step method)
```

13

```
	 Simultaneous Tests for General Linear Hypotheses

Fit: glmer(formula = cbind(X1, X2) ~ spec + (1 | GT), data = flo, 
    family = binomial)

Linear Hypotheses:
                    Estimate Std. Error z value Pr(>|z|)   
arc - fou == 0  -0.2370     0.1278  -1.854  0.15111   
arc - mic == 0  -0.5316     0.1534  -3.465  0.00147 **
fou - mic == 0  -0.2946     0.1573  -1.873  0.14514   
---
Signif. codes:  0 ‘*’ 0.001 ‘’ 0.01 ‘*’ 0.05 ‘.’ 0.1 ‘ ’ 1
(Adjusted p values reported -- single-step method)
```

14

```
	 Simultaneous Tests for General Linear Hypotheses

Fit: glmer(formula = cbind(X1, X2) ~ spec + (1 | GT), data = dead, 
    family = binomial)

Linear Hypotheses:
                    Estimate Std. Error z value Pr(>|z|)   
arc - fou == 0  -0.1995     0.1915  -1.042  0.54653   
arc - mic == 0   0.5756     0.2477   2.323  0.05156 . 
fou - mic == 0   0.7751     0.2617   2.962  0.00811 **
---
Signif. codes:  0 ‘*’ 0.001 ‘’ 0.01 ‘*’ 0.05 ‘.’ 0.1 ‘ ’ 1
(Adjusted p values reported -- single-step method)
```

#### TRAIT DIFFERENCES IN THE COMMON GARDEN, POPULATION LEVEL

1

```
	 Simultaneous Tests for General Linear Hypotheses

Fit: lmer(formula = Fd ~ gerGT + (1 | Host), data = FdAll, REML = T)

Linear Hypotheses:
                  Estimate Std. Error t value Pr(>|t|)    
C2 - C1 == 0   9.0064     2.1083   4.272  < 0.001 ***
C2 - G1 == 0   7.4620     1.9491   3.828  0.00178 ** 
C2 - G2 == 0  -1.5780     1.8971  -0.832  0.95944    
C2 - H2 == 0   0.2626     3.2781   0.080  1.00000    
C2 - H1 == 0  -3.9661     2.2516  -1.761  0.47836    
C1 - G1 == 0  -1.5444     1.9627  -0.787  0.96801    
C1 - G2 == 0 -10.5845     1.8913  -5.596  < 0.001 ***
C1 - H2 == 0  -8.7438     3.2703  -2.674  0.07700 .  
C1 - H1 == 0 -12.9726     2.2273  -5.824  < 0.001 ***
G1 - G2 == 0  -9.0401     1.7181  -5.262  < 0.001 ***
G1 - H2 == 0  -7.1994     3.1658  -2.274  0.19673    
G1 - H1 == 0 -11.4282     2.0885  -5.472  < 0.001 ***
G2 - H2 == 0   1.8406     3.1534   0.584  0.99161    
```

```
G2 - H1 == 0  -2.3881     2.0532  -1.163  0.84753
H2 - H1 == 0  -4.2287     3.3292  -1.270  0.79285
---
Signif. codes:  0 ‘*’ 0.001 ‘’ 0.01 ‘*’ 0.05 ‘.’ 0.1 ‘ ’ 1
(Adjusted p values reported -- single-step method)
```

2

```
	 Simultaneous Tests for General Linear Hypotheses

Fit: lmer(formula = Flower ~ gerGT + (1 | Host), data = FdAll, REML = T)

Linear Hypotheses:
                    Estimate Std. Error t value Pr(>|t|)
C2 - C1 == 0   3.1182     1.8338   1.700   0.5189
C2 - G1 == 0   5.6676     1.6953   3.343   0.0104 *
C2 - G2 == 0  -0.8171     1.6499  -0.495   0.9961
C2 - H2 == 0  -2.9667     2.8516  -1.040   0.8994
C2 - H1 == 0   1.9125     1.9583   0.977   0.9215
C1 - G1 == 0   2.5494     1.7073   1.493   0.6573
C1 - G2 == 0  -3.9353     1.6450  -2.392   0.1520
C1 - H2 == 0  -6.0849     2.8450  -2.139   0.2574
C1 - H1 == 0  -1.2057     1.9372  -0.622   0.9887
G1 - G2 == 0  -6.4847     1.4944  -4.339   <0.001 ***
G1 - H2 == 0  -8.6343     2.7539  -3.135   0.0206 *
G1 - H1 == 0  -3.7551     1.8166  -2.067   0.2943
G2 - H2 == 0  -2.1496     2.7434  -0.784   0.9686
G2 - H1 == 0   2.7296     1.7859   1.528   0.6342
H2 - H1 == 0   4.8792     2.8959   1.685   0.5295
---
Signif. codes:  0 ‘*’ 0.001 ‘’ 0.01 ‘*’ 0.05 ‘.’ 0.1 ‘ ’ 1
(Adjusted p values reported -- single-step method)
```

3

```
	 Simultaneous Tests for General Linear Hypotheses

Fit: lmer(formula = log(Plant.height) ~ gerGT + Transplant + (1 | 
       Host), data = FdAll, REML = T)

Linear Hypotheses:
                     Estimate Std. Error t value Pr(>|t|)
C2 - C1 == 0  0.065164   0.059561   1.094    0.878
C2 - G1 == 0 -0.224798   0.054823  -4.100   <0.001 ***
C2 - G2 == 0 -0.007977   0.053325  -0.150    1.000
C2 - H2 == 0 -0.068418   0.092264  -0.742    0.975
C2 - H1 == 0  0.319884   0.063568   5.032   <0.001 ***
C1 - G1 == 0 -0.289961   0.055350  -5.239   <0.001 ***
C1 - G2 == 0 -0.073140   0.053584  -1.365    0.738
C1 - H2 == 0 -0.133582   0.092052  -1.451    0.685
C1 - H1 == 0  0.254721   0.063713   3.998   <0.001 ***
G1 - G2 == 0  0.216821   0.048379   4.482   <0.001 ***
G1 - H2 == 0  0.156380   0.089074   1.756    0.482
G1 - H1 == 0  0.544682   0.059228   9.196   <0.001 ***
G2 - H2 == 0 -0.060441   0.088838  -0.680    0.983
G2 - H1 == 0  0.327861   0.057956   5.657   <0.001 ***
H2 - H1 == 0  0.388302   0.094090   4.127   <0.001 ***
---
Signif. codes:  0 ‘*’ 0.001 ‘’ 0.01 ‘*’ 0.05 ‘.’ 0.1 ‘ ’ 1
(Adjusted p values reported -- single-step method)
```

4

```
	 Simultaneous Tests for General Linear Hypotheses
```

```
Fit: lmer(formula = log(Height) ~ gerGT + Transplant + (1 | Host),
      data = FdAll, REML = T)

Linear Hypotheses:
                     Estimate Std. Error t value Pr(>|t|)
C2 - C1 == 0  0.03983    0.07882   0.505   0.9957
C2 - G1 == 0 -0.40317    0.07203  -5.597   <0.001 ***
C2 - G2 == 0 -0.06212    0.06960  -0.893   0.9452
C2 - H2 == 0 -0.07277    0.12512  -0.582   0.9917
C2 - H1 == 0  0.27149    0.08682   3.127   0.0209 *
C1 - G1 == 0 -0.44300    0.07390  -5.994   <0.001 ***
C1 - G2 == 0 -0.10195    0.07082  -1.439   0.6909
C1 - H2 == 0 -0.11259    0.12547  -0.897   0.9439
C1 - H1 == 0  0.23166    0.08753   2.647   0.0819 .
G1 - G2 == 0  0.34105    0.06360   5.362   <0.001 ***
G1 - H2 == 0  0.33040    0.12128   2.724   0.0670 .
G1 - H1 == 0  0.67466    0.08165   8.262   <0.001 ***
G2 - H2 == 0 -0.01064    0.12071  -0.088   1.0000
G2 - H1 == 0  0.33361    0.07959   4.192   <0.001 ***
H2 - H1 == 0  0.34425    0.12904   2.668   0.0778 .
---
Signif. codes:  0 ‘*’ 0.001 ‘’ 0.01 ‘*’ 0.05 ‘.’ 0.1 ‘ ’ 1
(Adjusted p values reported -- single-step method)
```

5

```
        Simultaneous Tests for General Linear Hypotheses

Fit: lmer(formula = log(1 + Reproductive.nodes) ~ gerGT + Transplant +
      (1 | Host), data = FdAll, REML = T)

Linear Hypotheses:
                     Estimate Std. Error t value Pr(>|t|)
C2 - C1 == 0  0.32497    0.09775   3.325  0.01101 *
C2 - G1 == 0  0.20349    0.08911   2.284  0.19146
C2 - G2 == 0  0.29909    0.08629   3.466  0.00684 **
C2 - H2 == 0  0.56422    0.15708   3.592  0.00426 **
C2 - H1 == 0  0.74632    0.11264   6.626  < 0.001 ***
C1 - G1 == 0 -0.12148    0.09166  -1.325  0.76016
C1 - G2 == 0 -0.02588    0.08804  -0.294  0.99968
C1 - H2 == 0  0.23925    0.15712   1.523  0.63599
C1 - H1 == 0  0.42135    0.11301   3.728  0.00269 **
G1 - G2 == 0  0.09560    0.07877   1.214  0.82148
G1 - H2 == 0  0.36073    0.15245   2.366  0.16024
G1 - H1 == 0  0.54283    0.10664   5.090  < 0.001 ***
G2 - H2 == 0  0.26514    0.15178   1.747  0.48593
G2 - H1 == 0  0.44723    0.10442   4.283  < 0.001 ***
H2 - H1 == 0  0.18210    0.16458   1.106  0.87198
---
Signif. codes:  0 ‘*’ 0.001 ‘’ 0.01 ‘*’ 0.05 ‘.’ 0.1 ‘ ’ 1
(Adjusted p values reported -- single-step method)
```

6

```
        Simultaneous Tests for General Linear Hypotheses

Fit: lmer(formula = Corolla.length ~ gerGT + (1 | Host), data = FdAll,
      REML = T)

Linear Hypotheses:
                      Estimate Std. Error t value Pr(>|t|)
C2 - C1 == 0  0.882641   0.091907   9.604   <0.001 ***
C2 - G1 == 0 -0.061978   0.084967  -0.729    0.977
C2 - G2 == 0 -0.009968   0.082697  -0.121    1.000
C2 - H2 == 0  1.908218   0.142909  13.353   <0.001 ***
C2 - H1 == 0  1.584933   0.098152  16.148   <0.001 ***
```

```
C1 - G1 == 0 -0.944619   0.085562 -11.040   <0.001 ***
C1 - G2 == 0 -0.892609   0.082448 -10.826   <0.001 ***
C1 - H2 == 0  1.025577   0.142574   7.193   <0.001 ***
C1 - H1 == 0  0.702292   0.097094   7.233   <0.001 ***
G1 - G2 == 0  0.052010   0.074899   0.694    0.982
G1 - H2 == 0  1.970196   0.138014  14.275   <0.001 ***
G1 - H1 == 0  1.646911   0.091046  18.089   <0.001 ***
G2 - H2 == 0  1.918186   0.137478  13.953   <0.001 ***
G2 - H1 == 0  1.594901   0.089506  17.819   <0.001 ***
H2 - H1 == 0 -0.323285   0.145136  -2.227    0.216
---
Signif. codes:  0 ‘*’ 0.001 ‘’ 0.01 ‘*’ 0.05 ‘.’ 0.1 ‘ ’ 1
(Adjusted p values reported -- single-step method)
```

7

```
        Simultaneous Tests for General Linear Hypotheses

Fit: lmer(formula = Nodes.to.flower ~ gerGT + (1 | Host), data = FdAll,
       REML = T)

Linear Hypotheses:
                     Estimate Std. Error t value Pr(>|t|)
C2 - C1 == 0   0.6138     0.1541   3.984   <0.001 ***
C2 - G1 == 0   1.5587     0.1424  10.946   <0.001 ***
C2 - G2 == 0   0.1655     0.1397   1.185   0.8373
C2 - H2 == 0   1.1529     0.2369   4.866   <0.001 ***
C2 - H1 == 0   1.6124     0.1641   9.827   <0.001 ***
C1 - G1 == 0   0.9449     0.1406   6.719   <0.001 ***
C1 - G2 == 0  -0.4483     0.1379  -3.251   0.0143 *
C1 - H2 == 0   0.5391     0.2359   2.286   0.1923
C1 - H1 == 0   0.9986     0.1625   6.144   <0.001 ***
G1 - G2 == 0  -1.3932     0.1247 -11.169   <0.001 ***
G1 - H2 == 0  -0.4058     0.2284  -1.776   0.4687
G1 - H1 == 0   0.0537     0.1515   0.354   0.9992
G2 - H2 == 0   0.9874     0.2268   4.355   <0.001 ***
G2 - H1 == 0   1.4469     0.1490   9.710   <0.001 ***
H2 - H1 == 0   0.4595     0.2425   1.895   0.3935
---
Signif. codes:  0 ‘*’ 0.001 ‘’ 0.01 ‘*’ 0.05 ‘.’ 0.1 ‘ ’ 1
(Adjusted p values reported -- single-step method)
```

8

```
        Simultaneous Tests for General Linear Hypotheses

Fit: lmer(formula = log(lir) ~ gerGT + (1 | Host), data = FdAll, REML = T)

Linear Hypotheses:
                     Estimate Std. Error t value Pr(>|t|)
C2 - C1 == 0  0.57830    0.07344   7.875  < 0.001 ***
C2 - G1 == 0  0.85583    0.06789  12.606  < 0.001 ***
C2 - G2 == 0  0.33111    0.06609   5.010  < 0.001 ***
C2 - H2 == 0  1.22982    0.11416  10.773  < 0.001 ***
C2 - H1 == 0  0.84126    0.07843  10.727  < 0.001 ***
C1 - G1 == 0  0.27753    0.06835   4.061  < 0.001 ***
C1 - G2 == 0 -0.24719    0.06588  -3.752  0.00234 **
C1 - H2 == 0  0.65151    0.11386   5.722  < 0.001 ***
C1 - H1 == 0  0.26296    0.07758   3.389  0.00889 **
G1 - G2 == 0 -0.52472    0.05984  -8.769  < 0.001 ***
G1 - H2 == 0  0.37398    0.11023   3.393  0.00876 **
G1 - H1 == 0 -0.01457    0.07274  -0.200  0.99995
G2 - H2 == 0  0.89871    0.10978   8.186  < 0.001 ***
G2 - H1 == 0  0.51015    0.07151   7.134  < 0.001 ***
H2 - H1 == 0 -0.38855    0.11594  -3.351  0.01020 *
---
Signif. codes:  0 ‘*’ 0.001 ‘’ 0.01 ‘*’ 0.05 ‘.’ 0.1 ‘ ’ 1
```

```
(Adjusted p values reported -- single-step method)
```

9

```
	 Simultaneous Tests for General Linear Hypotheses

Fit: lmer(formula = log(Number.of.leaf.teeth..exc..tip.) ~ gerGT + 
    Transplant + (1 | Host), data = FdAll)

Linear Hypotheses:
               Estimate Std. Error t value Pr(>|t|)    
C2 - C1 == 0  0.44314    0.04349  10.189  < 0.001 ***
C2 - G1 == 0  0.28003    0.04003   6.995  < 0.001 ***
C2 - G2 == 0  0.21931    0.03894   5.632  < 0.001 ***
C2 - H2 == 0  0.19283    0.06736   2.863  0.04608 *  
C2 - H1 == 0  0.54739    0.04642  11.793  < 0.001 ***
C1 - G1 == 0 -0.16312    0.04041  -4.036  < 0.001 ***
C1 - G2 == 0 -0.22384    0.03913  -5.721  < 0.001 ***
C1 - H2 == 0 -0.25032    0.06720  -3.725  0.00261 ** 
C1 - H1 == 0  0.10424    0.04652   2.241  0.21074    
G1 - G2 == 0 -0.06072    0.03532  -1.719  0.50654    
G1 - H2 == 0 -0.08720    0.06502  -1.341  0.75245    
G1 - H1 == 0  0.26736    0.04324   6.183  < 0.001 ***
G2 - H2 == 0 -0.02648    0.06485  -0.408  0.99845    
G2 - H1 == 0  0.32808    0.04232   7.753  < 0.001 ***
H2 - H1 == 0  0.35456    0.06869   5.162  < 0.001 ***
---
Signif. codes:  0 '*' 0.001 '' 0.01 '*' 0.05 '.' 0.1 ' ' 1
(Adjusted p values reported -- single-step method)
```

10

```
	 Simultaneous Tests for General Linear Hypotheses

Fit: lmer(formula = Capsule.width ~ gerGT + (1 | Host), data = FdAll, 
    REML = T)

Linear Hypotheses:
               Estimate Std. Error t value Pr(>|t|)    
C2 - C1 == 0 -0.10501    0.06424  -1.635    0.559    
C2 - G1 == 0 -0.30154    0.05714  -5.277   <0.001 ***
C2 - G2 == 0 -0.03239    0.05679  -0.570    0.992    
C2 - H2 == 0  0.45742    0.10756   4.253   <0.001 ***
C2 - H1 == 0  0.63594    0.08252   7.707   <0.001 ***
C1 - G1 == 0 -0.19653    0.06083  -3.231    0.015 *  
C1 - G2 == 0  0.07262    0.05989   1.212    0.821    
C1 - H2 == 0  0.56243    0.10802   5.207   <0.001 ***
C1 - H1 == 0  0.74095    0.08183   9.054   <0.001 ***
G1 - G2 == 0  0.26915    0.05227   5.149   <0.001 ***
G1 - H2 == 0  0.75896    0.10532   7.206   <0.001 ***
G1 - H1 == 0  0.93748    0.07939  11.809   <0.001 ***
G2 - H2 == 0  0.48981    0.10503   4.663   <0.001 ***
G2 - H1 == 0  0.66833    0.07942   8.415   <0.001 ***
H2 - H1 == 0  0.17852    0.11704   1.525    0.632    
---
Signif. codes:  0 '*' 0.001 '' 0.01 '*' 0.05 '.' 0.1 ' ' 1
(Adjusted p values reported -- single-step method)
```

11

```
	 Simultaneous Tests for General Linear Hypotheses

Fit: lmer(formula = Capsule.Height ~ gerGT + (1 | Host), data = FdAll, 
    REML = T)
```

```
Linear Hypotheses:
                          Estimate Std. Error t value Pr(>|t|)
C2 - C1 == 0  0.305012   0.151896   2.008  0.32285
C2 - G1 == 0 -0.581231   0.135065  -4.303  < 0.001 ***
C2 - G2 == 0 -0.589118   0.134242  -4.388  < 0.001 ***
C2 - H2 == 0  1.220697   0.254288   4.800  < 0.001 ***
C2 - H1 == 0  1.007755   0.195165   5.164  < 0.001 ***
C1 - G1 == 0 -0.886243   0.143848  -6.161  < 0.001 ***
C1 - G2 == 0 -0.894130   0.141612  -6.314  < 0.001 ***
C1 - H2 == 0  0.915684   0.255408   3.585  0.00448 **
C1 - H1 == 0  0.702743   0.193525   3.631  0.00382 **
G1 - G2 == 0 -0.007887   0.123574  -0.064  1.00000
G1 - H2 == 0  1.801928   0.248991   7.237  < 0.001 ***
G1 - H1 == 0  1.588987   0.187762   8.463  < 0.001 ***
G2 - H2 == 0  1.809815   0.248349   7.287  < 0.001 ***
G2 - H1 == 0  1.596874   0.187872   8.500  < 0.001 ***
H2 - H1 == 0 -0.212941   0.276651  -0.770  0.97031
---
Signif. codes:  0 ‘*’ 0.001 ‘’ 0.01 ‘*’ 0.05 ‘.’ 0.1 ‘ ’ 1
(Adjusted p values reported -- single-step method)
```

#### Supplemental text 2 – Models for *k*-mer spectra

A *k*-mer spectrum shows the distribution of counts of each individual *k*-mer (a small sequence word of length *k*) in a given data set. If the data set is generated from shotgun sequencing data of *one* individual, then the *k*-mer spectrum will reflect the genetic diversity in that individual's genome. The number of "peaks" in such a *k*-mer spectrum corresponds to the individual's ploidy level. A spectrum of polyploid has four peaks, which we label 1x, 2x, 3x, and 4x.

Genetic diversity is structured differently in autotetrapolyploids and allotetrapolyploids. While in autotetrapolyploids show tetrasomic inheritance, where recombination may happen between any pair of homologous chromosomes, allotetraploids contain two diverged (homoeologous) pairs of genomes. In allotetraploids, recombination may happen only within each homologous group, maintaining (and increasing over time) the genetic divergence between these groups. This causes different shapes of autotetraploid and allotetraploid *k*-mer spectra. For instance, allotetraploids should show a more pronounced 2x peak than autopolyploids, resulting from diverged alleles fixed in both homoeologous groups. The expectation for the shape of a *k*-mer spectrum can be obtained by combining the number of *k*-mers expected in each peak with a suitable distribution such as the negative binomial distribution as parameterised by Vurture et al. (2017). For autopolyploids, the (relative) number of *k*-mers expected in each peak can be obtained by using Ewens' (1972) sampling formula for the frequency of certain allelic configurations and by accounting for how many *k*-mers each configuration contributes to each peak of the *k*-mer spectrum. For allopolyploids, the homoeologous genomes can be treated like a pair of diverged sub-populations. The expectation for allelic configurations can then be obtained by the method of Lohse et al. (2011, 2016). The terms for the relative contributions to *k*-mer peaks in autotetraploids are:

$\frac{4\theta}{3+\theta}$ for the 1x peak,

$\frac{6\theta}{6+5\theta+\theta^2}$ for the 2x peak,

$\frac{8\theta}{6+11\theta+6\theta^2+\theta^3}$ for the 3x peak, and

$\frac{6}{6+11\theta+6\theta^2+\theta^3}$ for the 4x peak.

They all depend on $\theta$, the population-scaled mutation rate per *k*-mer ($\theta = 4N_e\mu$). From these formulae it can be shown that in an autotetraploid, the 2x peak can never contain more *k*-mers than the 1x peak: the inequality $\frac{4\theta}{3+\theta} < \frac{6\theta}{6+5\theta+\theta^2}$, which can be transformed into $6\theta + 17\theta^2 + 11\theta^3 + 2\theta^4 < 0$, cannot be true as long as the mutation rate is positive.

To compute the *k*-mer contributions in the allopolyploid model, an additional parameter $T$ is needed, the divergence time between the two homoeologous sub-genomes ($T$ is scaled in units of twice the effective population size $N_e$). The terms for the relative contributions in allopolyploids are:

$\frac{4e^{-3T\theta}\theta(-2e^{T(-2+\theta)}+e^{3T\theta}(2+\theta)^2(3+\theta)-2e^{2T\theta}(3+4\theta+\theta^2))}{(1+\theta)(2+\theta)^2(3+\theta)}$ for the 1x peak,

$\frac{2e^{-\frac{1}{2}T(4+5\theta)}(6e^{\frac{T\theta}{2}}\theta+e^{\frac{1}{2}T(4+5\theta)}(2+\theta)^2(3+\theta)+2e^{2T+\frac{3T\theta}{2}}(-6-8\theta+\theta^2+\theta^3))}{(1+\theta)(2+\theta)^2(3+\theta)}$ for the 2x peak,

$\frac{8e^{-T(3+4\theta)}\theta(-e^{T+2T\theta}+e^{3T(1+\theta)}(3+\theta))}{(1+\theta)(2+\theta)^2(3+\theta)}$ for the 3x peak, and

$\frac{2e^{-2T(1+\theta)}(\theta+2e^{T(2+\theta)}(3+\theta))}{(1+\theta)(2+\theta)^2(3+\theta)}$ for the 4x peak.

Ignoring any shared polymorphisms at the time of split, the divergence between the sub-genomes (per *k*-mer) can be computed as $\theta T$. This is reasonable, in particular for higher values of $T$, where most divergence will be due to alleles arising after the split of the sub-populations. A Mathematica notebook containing the code to derive the terms above is supplied in the Zenodo data set published alongside this paper.

In mixed maters, like many plant species, the estimate for $\theta$ obtained from one individual may be much lower than the diversity found in the population, as for $H_o$ and $H_e$. Because the sub-genome divergence time, $T$, is scaled by $N_e$, which itself depends on $\theta$, the estimate may not be meaningful as an estimate of the actual time of the split between the sub-genome progenitors. However, the average divergence per *k*-mer, $T\theta$, is not affected by this. When $T\theta$is divided by *k*, gives the average per-nucleotide sub-genome divergence.

### SUPPLEMENTAL TEXT 3 – GENOME PROFILING

Note: All analyses were run with *k*-mers of 27 nt length. For Sumdgeplot and Genomescope2.0, the haploid *k*-mer coverage was supplied as an input parameter.

### *Euphrasia anglica* (AN, diploid)

#### *GenomeScpe2.0*

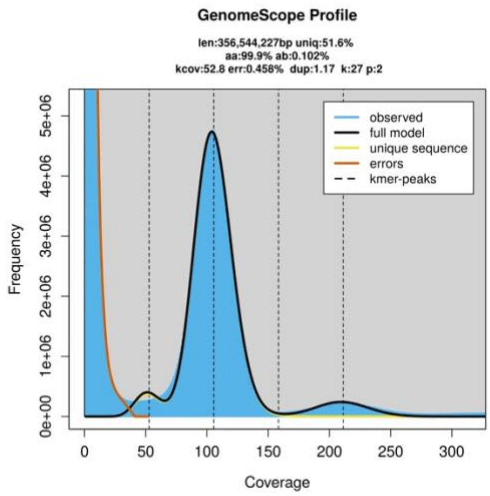

```
GenomeScope version 2.0
input file = /mnt/1TBSSD/Eukmers/numsE030
output directory = E030
p = 2
k = 27
initial kmercov estimate = 53

property                      min               max
Homozygous (aa)               99.8858%          99.911%
Heterozygous (ab)             0.0889549%        0.114179%
Genome Haploid Length         355,898,845 bp    356,544,227 bp
Genome Repeat Length          172,096,499 bp    172,408,576 bp
Genome Unique Length          183,802,346 bp    184,135,651 bp
Model Fit                     56.7642%          94.5677%
Read Error Rate               0.457507%         0.457507%
```

#### *Smudgeplot*

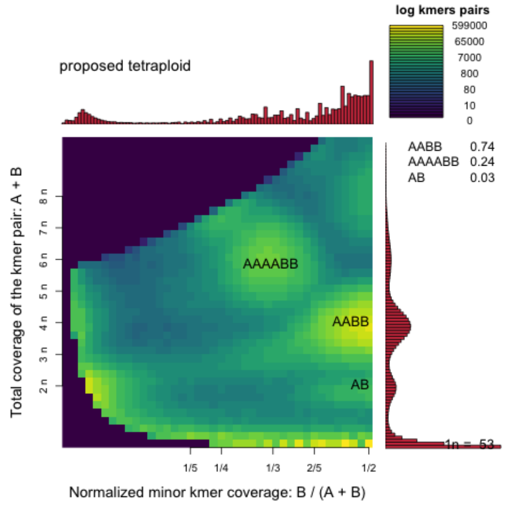

#### *Tetmer*

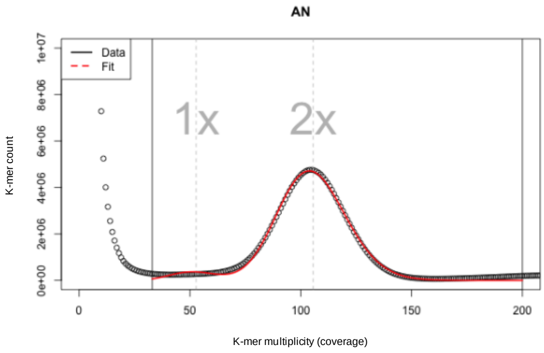

```
DIPLOID MODEL, AUTO FITTED
     haploid k-mer cov: 52.8
       per k-mer theta: 0.0273
hapl non-rep GS (Mbp): 185.3
     bias (peak width): 1.2

STARTING RANGES (MIN MAX)
     haploid k-mer cov: 19 59
log10 per k-mer theta: -2.75 0.6
hapl non-rep GS (Mbp): 20 2000
     bias (peak width): 0.1 3
              x range: 33 200
```

#### *EUPHRASIA VIGURSII* (VI, DIPLOID)

##### GENOMESCPE2.0

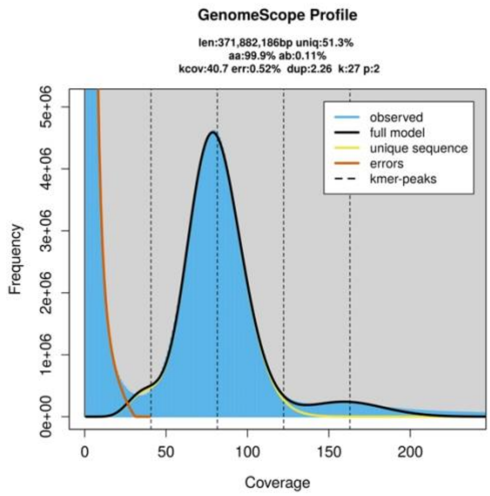

```
GenomeScope version 2.0
input file = /mnt/1TBSSD/Eukmers/numsE031
output directory = E031
p = 2
k = 27
initial kmercov estimate = 41

property                      min               max
Homozygous (aa)               99.8731%          99.9071%
Heterozygous (ab)             0.0928852%        0.126899%
Genome Haploid Length         370,710,233 bp    371,882,186 bp
Genome Repeat Length          180,417,372 bp    180,987,738 bp
Genome Unique Length          190,292,861 bp    190,894,447 bp
Model Fit                     56.9501%          96.4987%
Read Error Rate               0.519548%         0.519548%
```

##### SMUDGEPLOT

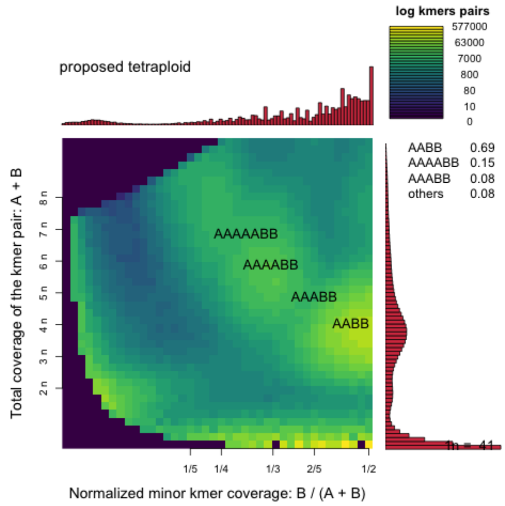

##### TETMER

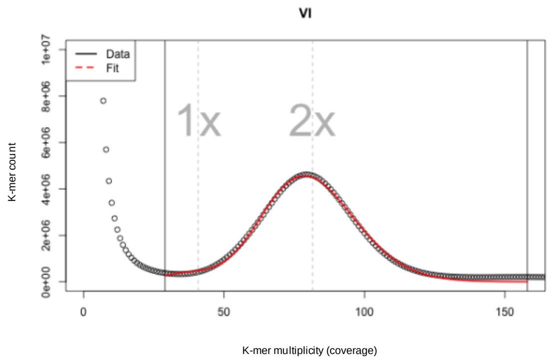

```
DIPLOID MODEL, AUTO FITTED
     haploid k-mer cov: 40.8
       per k-mer theta: 0.0294
hapl non-rep GS (Mbp): 192.6
     bias (peak width): 2.4

STARTING RANGES (MIN MAX)
     haploid k-mer cov: 10 55
log10 per k-mer theta: -2.4 0.6
hapl non-rep GS (Mbp): 20 2000
     bias (peak width): 0.1 3
              x range: 29 158
```

#### *EUPHRASIA RIVULARIS* (RI, DIPLOID)

##### GENOMESCPE2.0

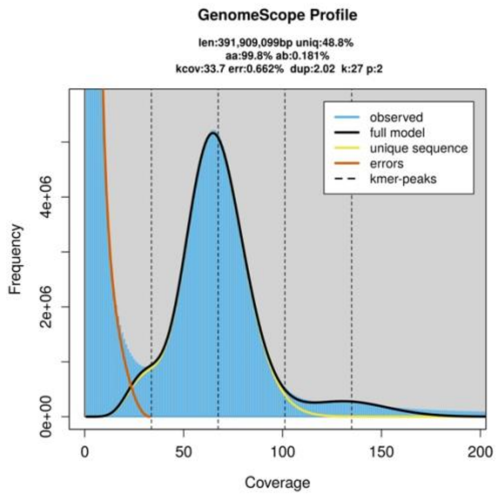

```
GenomeScope version 2.0
input file = /mnt/1TBSSD/Eukmers/numsE032
output directory = E032
p = 2
k = 27
initial kmercov estimate = 33

property                      min              max
Homozygous (aa)               99.7973%         99.8408%
Heterozygous (ab)             0.159239%        0.202729%
Genome Haploid Length         390,392,793 bp   391,909,099 bp
Genome Repeat Length          199,832,645 bp   200,608,806 bp
Genome Unique Length          190,560,148 bp   191,300,294 bp
Model Fit                     54.4646%         97.309%
Read Error Rate               0.66216%         0.66216%
```

##### SMUDGEPLOT

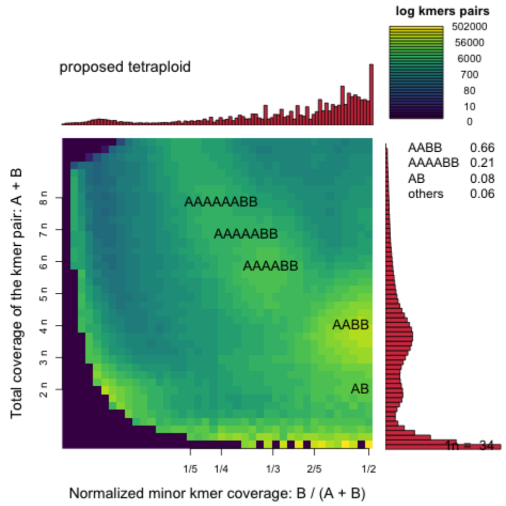

##### TETMER

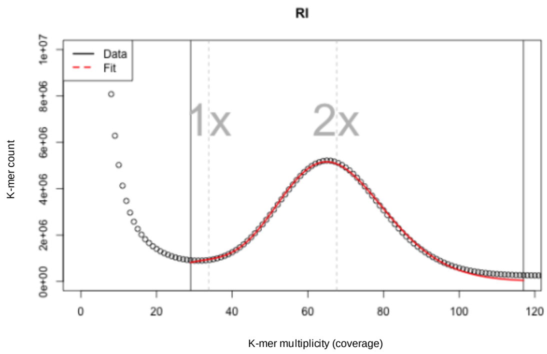

```
DIPLOID MODEL, AUTO FITTED
     haploid k-mer cov: 33.8
       per k-mer theta: 0.0542
hapl non-rep GS (Mbp): 193.1
     bias (peak width): 2.1

STARTING RANGES (MIN MAX)
     haploid k-mer cov: 10 41
log10 per k-mer theta: -2 0.6
hapl non-rep GS (Mbp): 20 2000
     bias (peak width): 0.1 3
              x range: 29 117
```

#### *EUPHRASIA ROSTKOVIANA* (RO, DIPLOID)

##### *GENOMESCPE2.0*

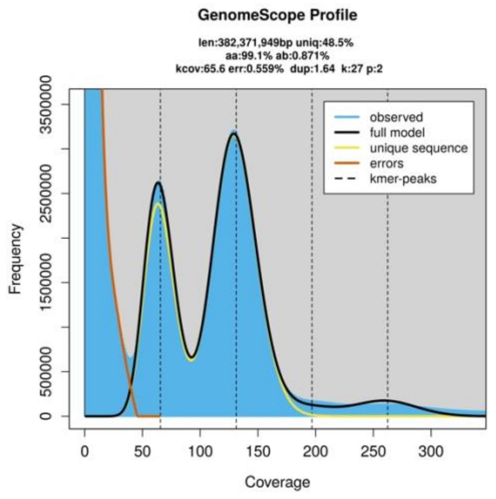

```
GenomeScope version 2.0
input file = /mnt/1TBSSD/Eukmers/numsE040
output directory = E040
p = 2
k = 27
initial kmercov estimate = 66

property                    min             max
Homozygous (aa)             99.1141%        99.1449%
Heterozygous (ab)           0.855141%       0.885937%
Genome Haploid Length       381,376,095 bp  382,371,949 bp
Genome Repeat Length        196,460,140 bp  196,973,139 bp
Genome Unique Length        184,915,956 bp  185,398,811 bp
Model Fit                   55.4459%        94.5013%
Read Error Rate             0.559427%       0.559427%
```

##### *SMUDGEPLOT*

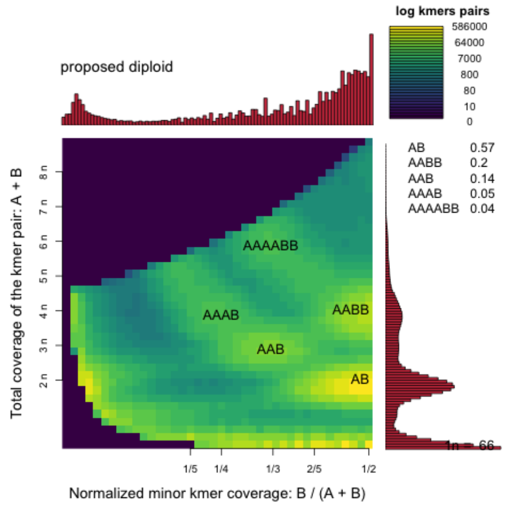

##### *TETMER*

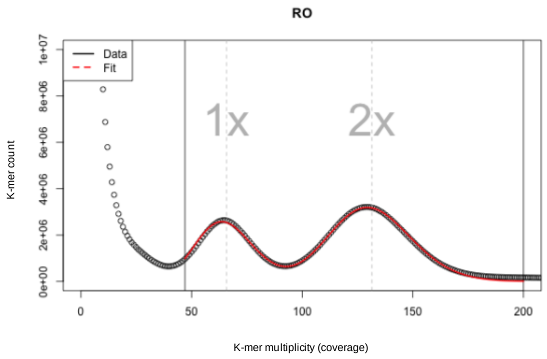

```
DIPLOID MODEL, AUTO FITTED
    haploid k-mer cov: 65.7
      per k-mer theta: 0.2841
hapl non-rep GS (Mbp): 189.5
    bias (peak width): 1.6

STARTING RANGES (MIN MAX)
    haploid k-mer cov: 10 85
log10 per k-mer theta: -2 0.6
hapl non-rep GS (Mbp): 20 2000
    bias (peak width): 0.1 3
              x range: 47 200
```

#### *Euphrasia arctica* (A0, tetraploid)

##### *GenomeScpe2.0*

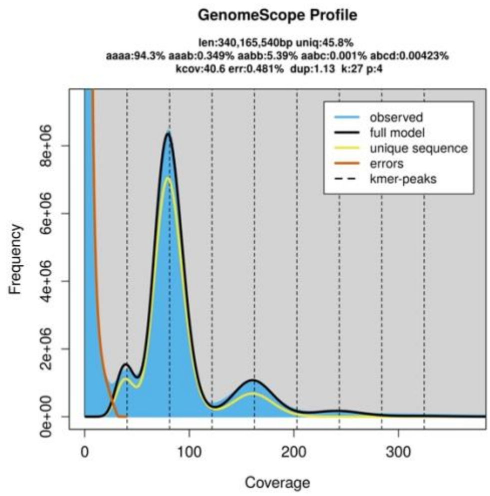

```
GenomeScope version 2.0
input file = /mnt/1TBSSD/Eukmers/numsE028
output directory = E028
p = 4
k = 27
initial kmercov estimate = 40

property                      min               max
Homozygous (aaaa)             92.4957%          94.7871%
Heterozygous (not aaaa)       5.21291%          7.50425%
aaab                          0%                1.04367%
aabb                          5.21291%          5.57398%
aabc                          0%                0.382878%
abcd                          0%                0.503721%
Genome Haploid Length         339,487,037 bp    340,165,540 bp
Genome Repeat Length          183,904,958 bp    184,272,513 bp
Genome Unique Length          155,582,079 bp    155,893,027 bp
Model Fit                     60.4511%          91.886%
Read Error Rate               0.480646%         0.480646%
```

##### *Smudgeplot*

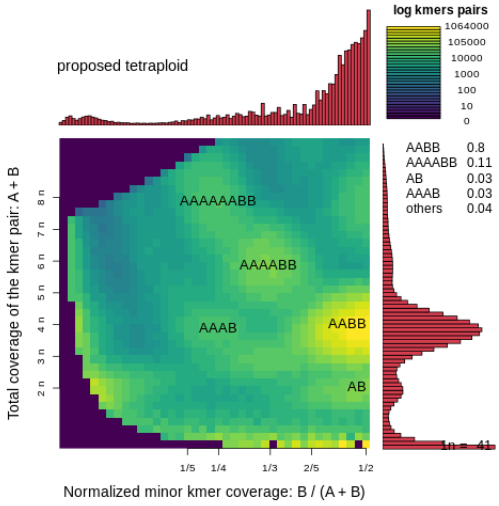

##### *Tetmer*

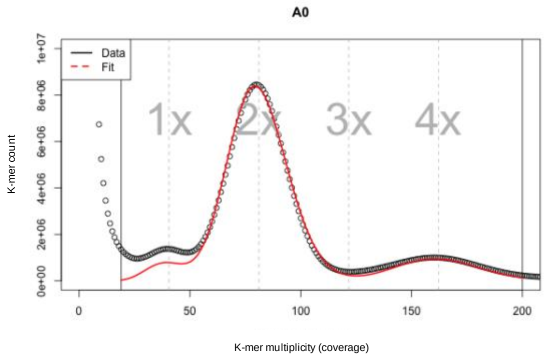

```
ALLOTETRAPLOID MODEL, AUTO FITTED
     haploid k-mer cov: 40.6
       per k-mer theta: 0.0279
                     T: 49.99
hapl non-rep GS (Mbp): 191.2
     bias (peak width): 1.2
      per k-mer diverg: 1.395

STARTING RANGES (MIN MAX)
     haploid k-mer cov: 21 53
log10 per k-mer theta: -2.95 0.6
                     T: 0.001 100
hapl non-rep GS (Mbp): 20 2000
     bias (peak width): 0.1 3
             x range: 19 200
```

### *EUPHRASIA ARCTICA* (A1, TETRAPLOID)

#### GENOMESCPE2.0

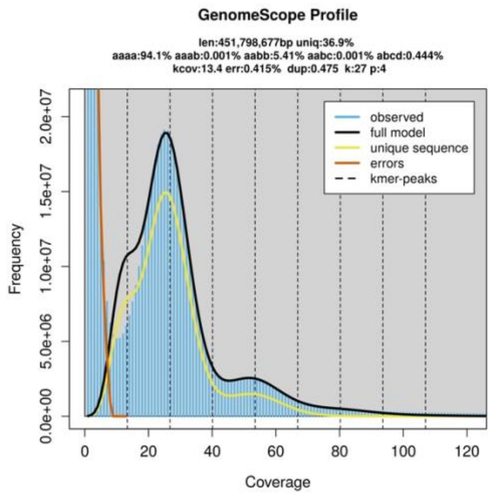

```
GenomeScope version 2.0
input file = /mnt/1TBSSD/Eukmers/numsE027
output directory = E027
p = 4
k = 27
initial kmercov estimate = 13

property                      min              max
Homozygous (aaaa)             91.5436%         95.0556%
Heterozygous (not aaaa)       4.94439%         8.45638%
aaab                          0%               0.790602%
aabb                          4.94439%         5.88006%
aabc                          0%               0.739464%
abcd                          0%               1.04626%
Genome Haploid Length         448,652,997 bp   451,798,677 bp
Genome Repeat Length          283,222,006 bp   285,207,786 bp
Genome Unique Length          165,430,991 bp   166,590,891 bp
Model Fit                     51.1329%         92.7128%
Read Error Rate               0.414595%        0.414595%
```

#### SMUDGEPLOT

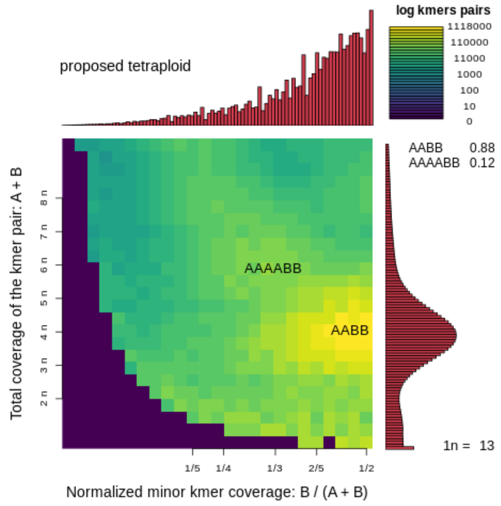

#### TETMER

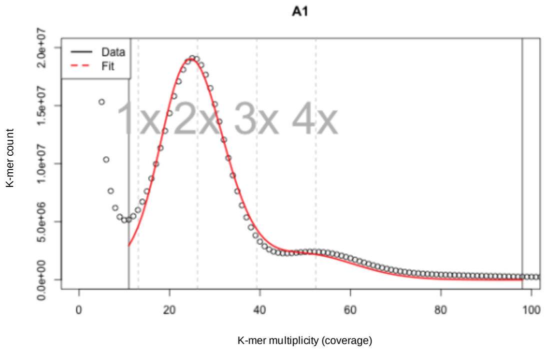

```
ALLOTETRAPLOID MODEL, AUTO FITTED
    haploid k-mer cov: 13.1
      per k-mer theta: 0.028
                    T: 50.01
hapl non-rep GS (Mbp): 224.3
    bias (peak width): 0.9
     per k-mer diverg: 1.399

STARTING RANGES (MIN MAX)
    haploid k-mer cov: 10 18
log10 per k-mer theta: -4 0.6
                    T: 0.001 100
hapl non-rep GS (Mbp): 20 2000
    bias (peak width): 0.1 3
              x range: 11 98
```

#### *EUPHRASIA ARCTICA* (A2, TETRAPLOID)

##### GENOMESCPE2.0

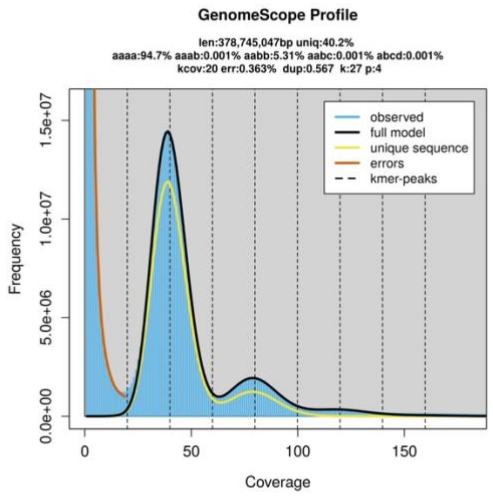

```
GenomeScope version 2.0
input file = /mnt/1TBSSD/Eukmers/numsE026
output directory = E026
p = 4
k = 27
initial kmercov estimate = 20

property                    min             max
Homozygous (aaaa)           92.7655%        94.8998%
Heterozygous (not aaaa)     5.10016%        7.23446%
aaab                        0%              0.759079%
aabb                        5.10016%        5.52219%
aabc                        0%              0.392766%
abcd                        0%              0.560428%
Genome Haploid Length       376,975,633 bp  378,745,047 bp
Genome Repeat Length        225,294,331 bp  226,351,798 bp
Genome Unique Length        151,681,302 bp  152,393,250 bp
Model Fit                   55.8344%        94.6734%
Read Error Rate             0.362751%       0.362751%
```

##### SMUDGEPLOT

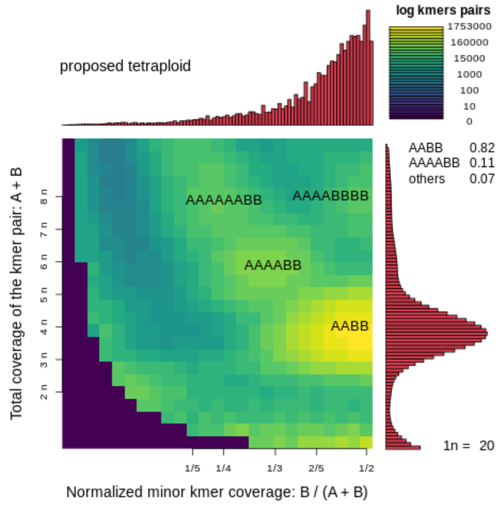

##### TETMER

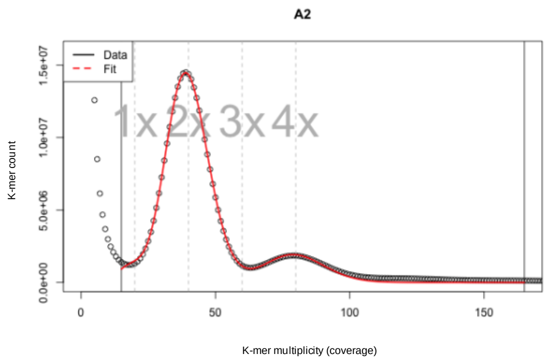

```
ALLOTETRAPLOID MODEL, AUTO FITTED
     haploid k-mer cov: 20
       per k-mer theta: 0.0244
                     T: 52.45
hapl non-rep GS (Mbp): 201.1
     bias (peak width): 0.5
      per k-mer diverg: 1.279

STARTING RANGES (MIN MAX)
     haploid k-mer cov: 10 34
log10 per k-mer theta: -3.2 0.6
                     T: 5 100
hapl non-rep GS (Mbp): 89 373
     bias (peak width): 0.1 3
             x range: 15 165
```

#### *Euphrasia arctica* (A3, tetraploid)

##### GenomeScpe2.0

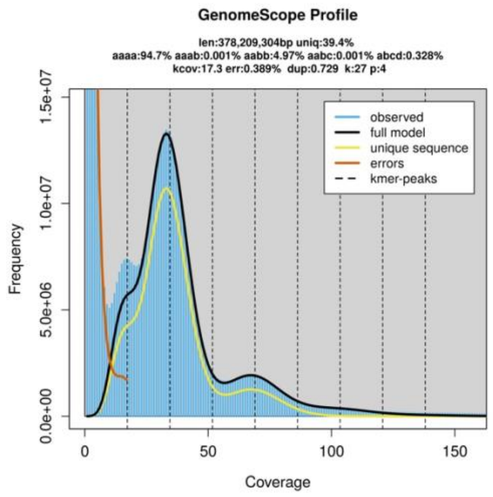

```
GenomeScope version 2.0
input file = /mnt/1TBSSD/Eukmers/numsE036
output directory = E036
p = 4
k = 27
initial kmercov estimate = 17

property                      min              max
Homozygous (aaaa)             92.8383%         95.1968%
Heterozygous (not aaaa)       4.80318%         7.16166%
aaab                          0%               0.584296%
aabb                          4.80318%         5.13048%
aabc                          0%               0.606882%
abcd                          0%               0.839998%
Genome Haploid Length         376,114,344 bp   378,209,304 bp
Genome Repeat Length          227,949,510 bp   229,219,190 bp
Genome Unique Length          148,164,835 bp   148,990,114 bp
Model Fit                     55.5124%         94.4969%
Read Error Rate               0.388542%        0.388542%
```

##### Smudgeplot

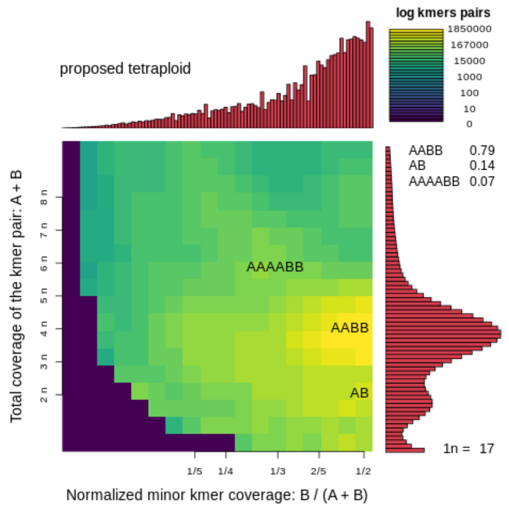

##### Tetmer

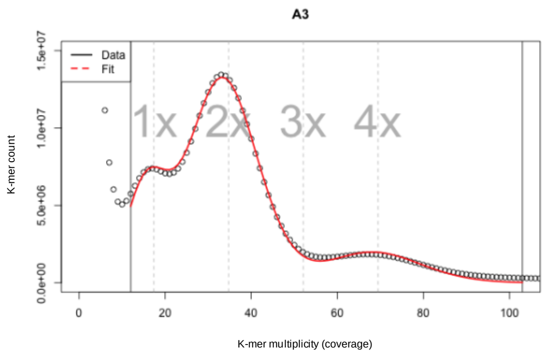

```
ALLOTETRAPLOID MODEL, AUTO FITTED
     haploid k-mer cov: 17.4
        per k-mer theta: 0.1507
                      T: 7.28
hapl non-rep GS (Mbp): 213.1
     bias (peak width): 0.7
       per k-mer diverg: 1.097

STARTING RANGES (MIN MAX)
     haploid k-mer cov: 10 24
log10 per k-mer theta: -2 0.8
                      T: 0.001 100
hapl non-rep GS (Mbp): 20 400
     bias (peak width): 0.01 0.96
              x range: 12 103
```

#### *EUPHRASIA FOULAENSIS* (F1, TETRAPLOID)

##### GENOMESCPE2.0

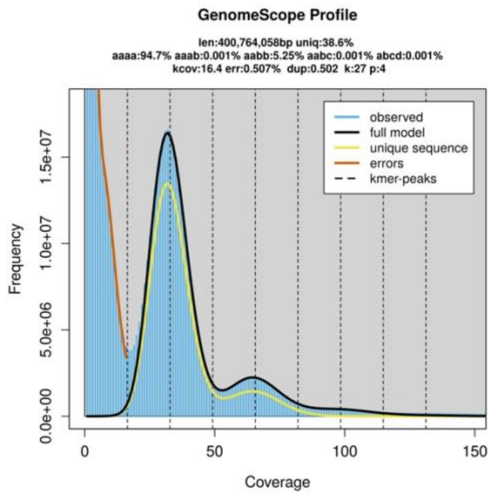

```
GenomeScope version 2.0
input file = /mnt/1TBSSD/Eukmers/numsE025
output directory = E025
p = 4
k = 27
initial kmercov estimate = 16

property                      min              max
Homozygous (aaaa)             91.7601%         94.9911%
Heterozygous (not aaaa)       5.00888%         8.23991%
aaab                          0%               1.08841%
aabb                          5.00888%         5.48638%
aabc                          0%               0.678665%
abcd                          0%               0.986456%
Genome Haploid Length         395,889,466 bp   400,764,058 bp
Genome Repeat Length          243,095,268 bp   246,088,503 bp
Genome Unique Length          152,794,198 bp   154,675,555 bp
Model Fit                     53.763%          93.1246%
Read Error Rate               0.5072%          0.5072%
```

##### SMUDGEPLOT

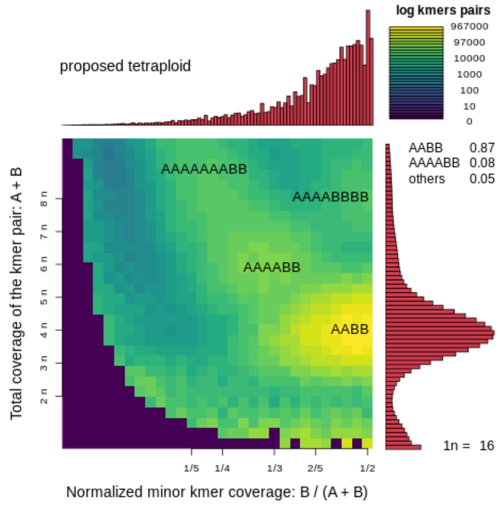

##### TETMER

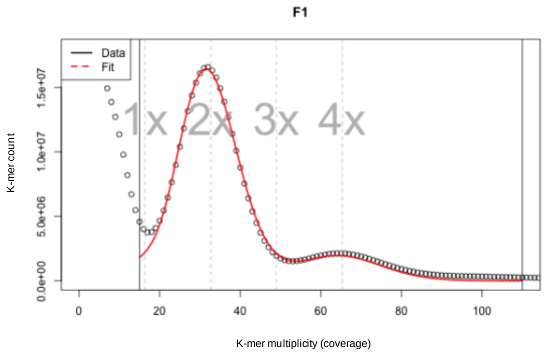

```
ALLOTETRAPLOID MODEL, AUTO FITTED
    haploid k-mer cov: 16.3
      per k-mer theta: 0.0244
                    T: 55.55
hapl non-rep GS (Mbp): 204.1
    bias (peak width): 0.6
     per k-mer diverg: 1.357

STARTING RANGES (MIN MAX)
    haploid k-mer cov: 10 24
log10 per k-mer theta: -3.3 0.6
                    T: 0.001 100
hapl non-rep GS (Mbp): 61 317
    bias (peak width): 0.1 3
              x range: 15 110
```

#### *EUPHRASIA FOULAENSIS* (F2, TETRAPLOID)

##### GENOMESCPE2.0

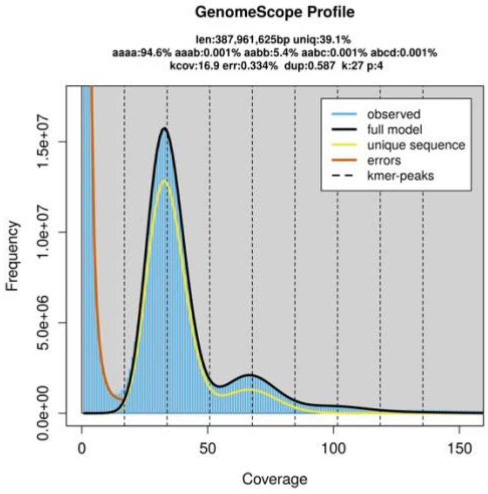

```
GenomeScope version 2.0
input file = /mnt/1TBSSD/Eukmers/numsE024
output directory = E024
p = 4
k = 27
initial kmercov estimate = 16

property                      min              max
Homozygous (aaaa)             92.3039%         94.9094%
Heterozygous (not aaaa)       5.09063%         7.69614%
aaab                          0%               0.888598%
aabb                          5.09063%         5.70509%
aabc                          0%               0.428555%
abcd                          0%               0.673904%
Genome Haploid Length         384,406,674 bp   387,961,625 bp
Genome Repeat Length          234,199,161 bp   236,365,009 bp
Genome Unique Length          150,207,513 bp   151,596,616 bp
Model Fit                     55.237%          95.2557%
Read Error Rate               0.334242%        0.334242%
```

##### SMUDGEPLOT

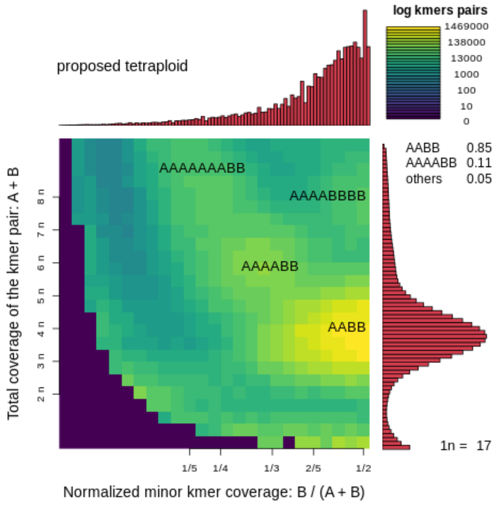

##### TETMER

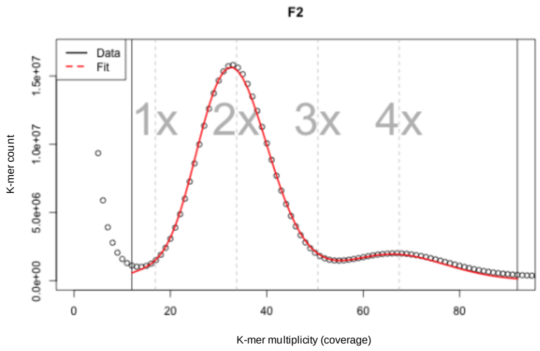

```
ALLOTETRAPLOID MODEL, AUTO FITTED
     haploid k-mer cov: 16.9
       per k-mer theta: 0.0133
                     T: 100
hapl non-rep GS (Mbp): 199.4
     bias (peak width): 0.7
      per k-mer diverg: 1.334

STARTING RANGES (MIN MAX)
     haploid k-mer cov: 11 21
log10 per k-mer theta: -3.7 0.2
                     T: 0.001 100
hapl non-rep GS (Mbp): 130 331
     bias (peak width): 0.1 1.36
              x range: 12 92
```

### *EUPHRASIA FOULAENSIS* (F3, TETRAPLOID)

#### GENOMESCPE2.0

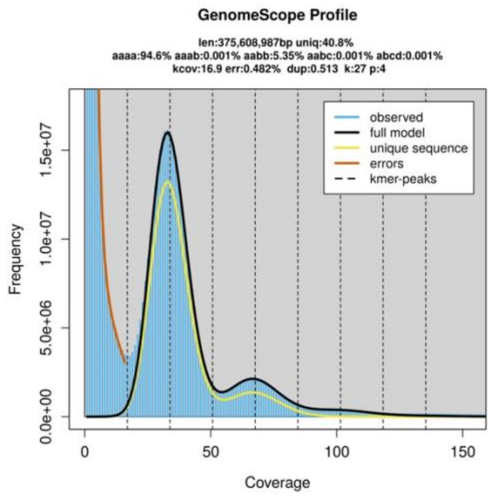

```
GenomeScope version 2.0
input file = /mnt/1TBSSD/Eukmers/numsE038
output directory = E038
p = 4
k = 27
initial kmercov estimate = 16

property                      min              max
Homozygous (aaaa)             91.8757%         94.8902%
Heterozygous (not aaaa)       5.10979%         8.12434%
aaab                          0%               1.0092%
aabb                          5.10979%         5.59446%
aabc                          0%               0.577994%
abcd                          0%               0.942694%
Genome Haploid Length         371,448,295 bp   375,608,987 bp
Genome Repeat Length          220,027,222 bp   222,491,806 bp
Genome Unique Length          151,421,073 bp   153,117,181 bp
Model Fit                     56.4736%         93.702%
Read Error Rate               0.482457%        0.482457%
```

#### SMUDGEPLOT

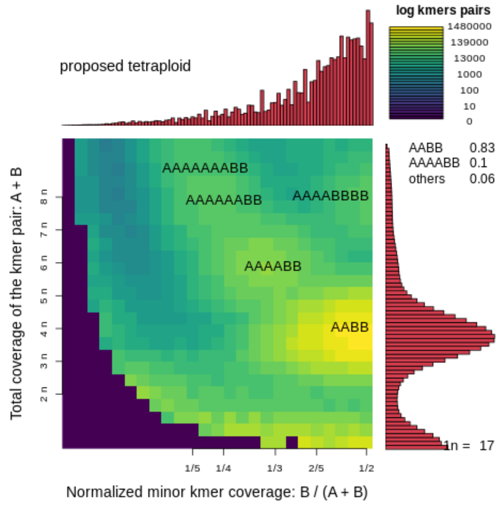

#### TETMER

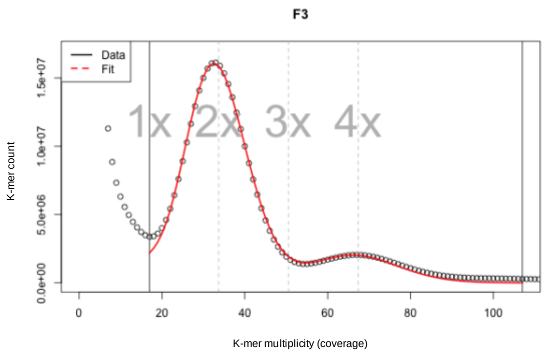

```
ALLOTETRAPLOID MODEL, AUTO FITTED
     haploid k-mer cov: 16.8
       per k-mer theta: 0.0262
                     T: 50
hapl non-rep GS (Mbp): 205.1
     bias (peak width): 0.6
      per k-mer diverg: 1.312

STARTING RANGES (MIN MAX)
     haploid k-mer cov: 10 34
log10 per k-mer theta: -2 0.6
                     T: 0.001 100
hapl non-rep GS (Mbp): 20 2000
     bias (peak width): 0.1 3
              x range: 17 107
```

#### *EUPHRASIA FOULAENSIS* (F4, TETRAPLOID)

##### GENOMESCPE2.0

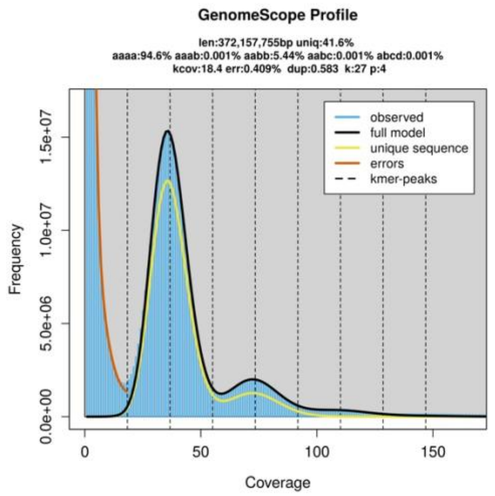

```
GenomeScope version 2.0
input file = /mnt/1TBSSD/Eukmers/numsE037
output directory = E037
p = 4
k = 27
initial kmercov estimate = 18

property                    min              max
Homozygous (aaaa)           92.3052%         94.815%
Heterozygous (not aaaa)     5.18496%         7.69476%
aaab                        0%               0.859625%
aabb                        5.18496%         5.70374%
aabc                        0%               0.416018%
abcd                        0%               0.715371%
Genome Haploid Length       369,079,264 bp   372,157,755 bp
Genome Repeat Length        215,414,425 bp   217,211,197 bp
Genome Unique Length        153,664,839 bp   154,946,558 bp
Model Fit                   57.621%          94.1044%
Read Error Rate             0.408706%        0.408706%
```

##### SMUDGEPLOT

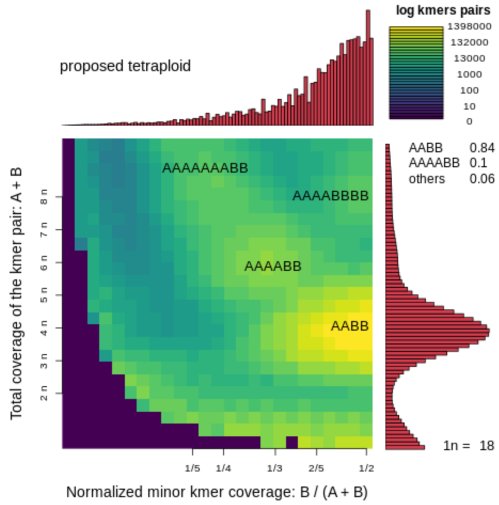

##### TETMER

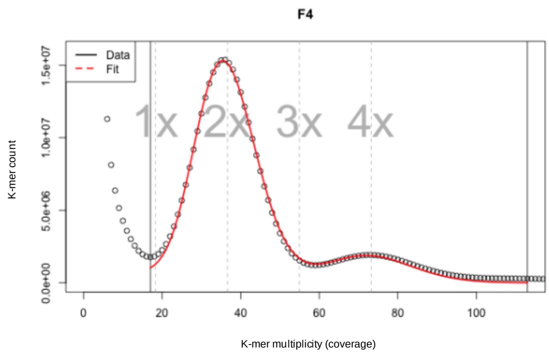

```
ALLOTETRAPLOID MODEL, AUTO FITTED
     haploid k-mer cov: 18.3
       per k-mer theta: 0.0134
                     T: 100
hapl non-rep GS (Mbp): 203.8
     bias (peak width): 0.7
      per k-mer diverg: 1.343

STARTING RANGES (MIN MAX)
     haploid k-mer cov: 17 31
log10 per k-mer theta: -2.65 0.6
                     T: 0.001 100
hapl non-rep GS (Mbp): 20 2000
     bias (peak width): 0.1 3
               x range: 17 113
```

### *Euphrasia micrantha* (M0, tetraploid)

#### GenomeScpe2.0

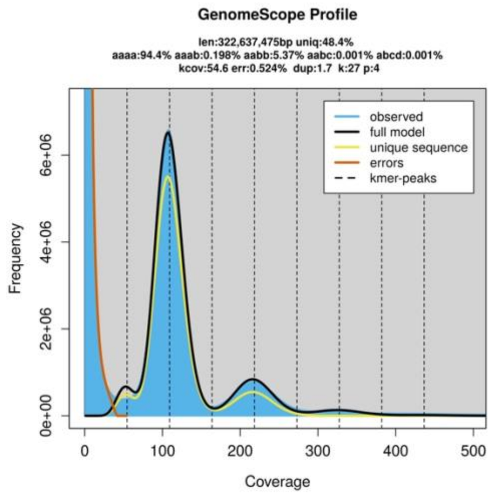

```
GenomeScope version 2.0
input file = /mnt/1TBSSD/Eukmers/numsE033
output directory = E033
p = 4
k = 27
initial kmercov estimate = 54

property                    min               max
Homozygous (aaaa)           92.7157%          94.7738%
Heterozygous (not aaaa)     5.22624%          7.28431%
aaab                        0%                0.851321%
aabb                        5.22624%          5.52321%
aabc                        0%                0.406022%
abcd                        0%                0.503752%
Genome Haploid Length       322,049,625 bp    322,637,475 bp
Genome Repeat Length        166,242,735 bp    166,546,184 bp
Genome Unique Length        155,806,890 bp    156,091,291 bp
Model Fit                   63.1628%          91.6415%
Read Error Rate             0.524025%         0.524025%
```

#### Smudgeplot

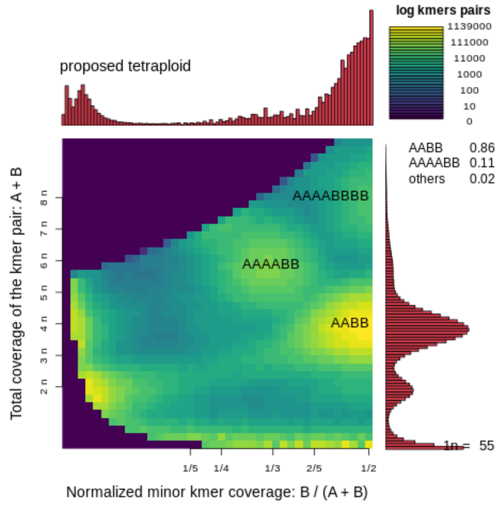

#### Tetmer

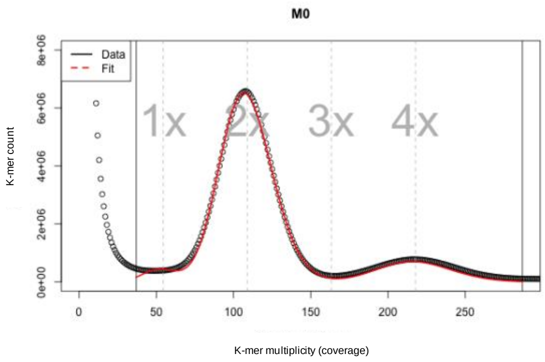

```
ALLOTETRAPLOID MODEL, AUTO FITTED
    haploid k-mer cov: 54.4
      per k-mer theta: 0.0205
                    T: 70.13
hapl non-rep GS (Mbp): 185.1
    bias (peak width): 1.7
     per k-mer diverg: 1.437

STARTING RANGES (MIN MAX)
    haploid k-mer cov: 34 63
log10 per k-mer theta: -3.85 -1.55
                    T: 0.001 100
hapl non-rep GS (Mbp): 117 345
```

```
                  bias (peak width): 0.1 3
                             x range: 37 287
```

#### *Euphrasia micrantha* (M1, tetraploid)

##### GenomeScpe2.0

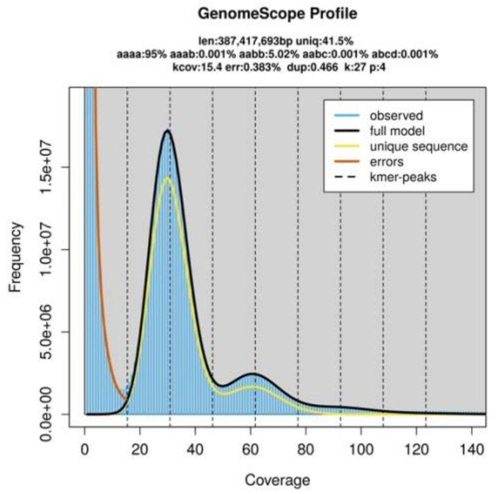

```
GenomeScope version 2.0
input file = /mnt/1TBSSD/Eukmers/numsE023
output directory = E023
p = 4
k = 27
initial kmercov estimate = 15

property                       min                 max
Homozygous (aaaa)              92.6876%            95.1105%
Heterozygous (not aaaa)        4.88955%            7.31239%
aaab                           0%                  0.732638%
aabb                           4.88955%            5.14633%
aabc                           0%                  0.651672%
abcd                           0%                  0.781753%
Genome Haploid Length          382,918,749 bp      387,417,693 bp
Genome Repeat Length           223,967,285 bp      226,598,695 bp
Genome Unique Length           158,951,464 bp      160,818,998 bp
Model Fit                      56.8672%            95.5142%
Read Error Rate                0.383123%           0.383123%
```

##### Smudgeplot

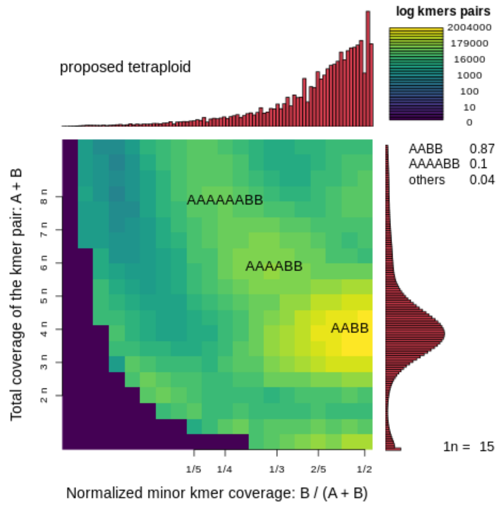

##### Tetmer

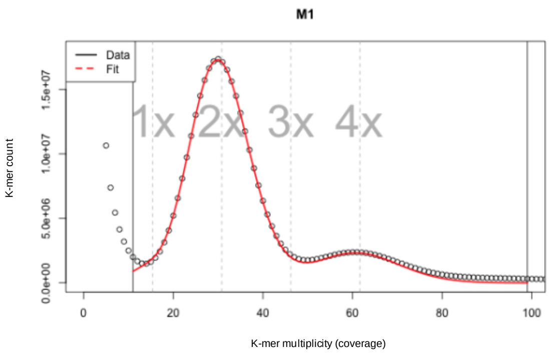

```
ALLOTETRAPLOID MODEL, AUTO FITTED
    haploid k-mer cov: 15.4
      per k-mer theta: 0.0196
                    T: 64.79
hapl non-rep GS (Mbp): 201.7
    bias (peak width): 0.4
     per k-mer diverg: 1.272

STARTING RANGES (MIN MAX)
    haploid k-mer cov: 10 18
log10 per k-mer theta: -4 0.6
                    T: 0.001 80
hapl non-rep GS (Mbp): 20 400
```

```
    bias (peak width): 0.1 1.66
             x range: 11 99
```

#### *Euphrasia micrantha* (M2, tetraploid)

##### GenomeScpe2.0

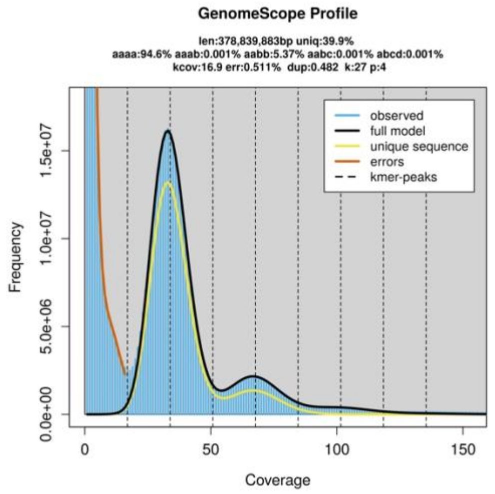

```
GenomeScope version 2.0
input file = /mnt/1TBSSD/Eukmers/numsE022
output directory = E022
p = 4
k = 27
initial kmercov estimate = 17

property                     min              max
Homozygous (aaaa)            91.796%          94.9132%
Heterozygous (not aaaa)      5.08677%         8.20403%
aaab                         0%               1.05657%
aabb                         5.08677%         5.64331%
aabc                         0%               0.583533%
abcd                         0%               0.920616%
Genome Haploid Length        374,837,102 bp   378,839,883 bp
Genome Repeat Length         225,133,438 bp   227,537,576 bp
Genome Unique Length         149,703,664 bp   151,302,307 bp
Model Fit                    56.0751%         94.4486%
Read Error Rate              0.510853%        0.510853%
```

##### Smudgeplot

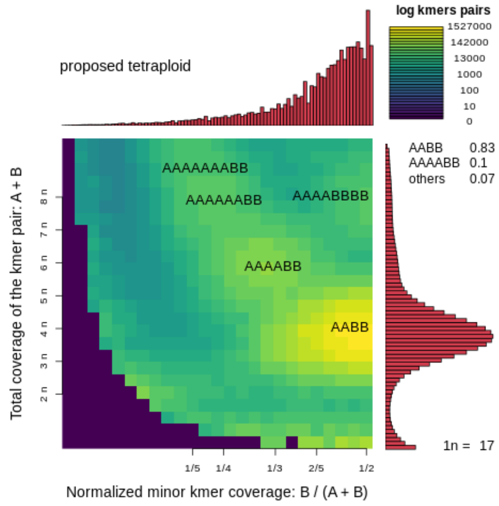

##### Tetmer

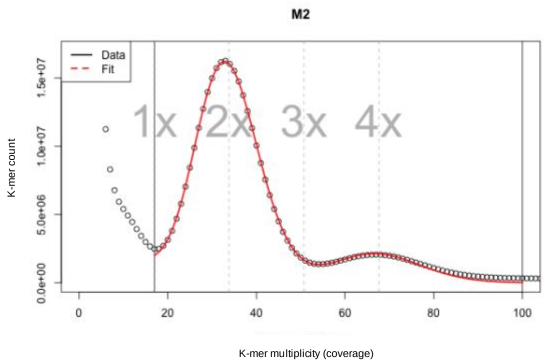

```
ALLOTETRAPLOID MODEL, AUTO FITTED
     haploid k-mer cov: 16.9
        per k-mer theta: 0.0256
                      T: 50.03
hapl non-rep GS (Mbp): 202.7
     bias (peak width): 0.5
      per k-mer diverg: 1.283

STARTING RANGES (MIN MAX)
     haploid k-mer cov: 15 24
log10 per k-mer theta: -3.75 0.6
```

```
                       T: 0.001 100
hapl non-rep GS (Mbp): 20 2000
    bias (peak width): 0.1 3
             x range: 17 100
```

#### *EUPHRASIA MICRANTHA* (M3, TETRAPLOID)

##### GENOMESCPE2.0

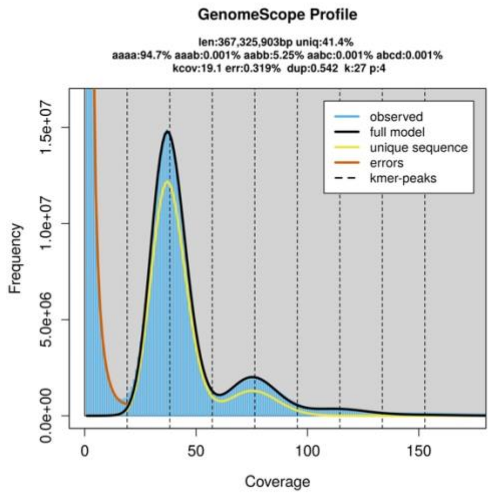

```
GenomeScope version 2.0
input file = /mnt/1TBSSD/Eukmers/numsE035
output directory = E035
p = 4
k = 27
initial kmercov estimate = 19

property                    min              max
Homozygous (aaaa)           92.8567%         94.9359%
Heterozygous (not aaaa)     5.06408%         7.14332%
aaab                        0%               0.7246%
aabb                        5.06408%         5.4417%
aabc                        0%               0.397073%
abcd                        0%               0.579947%
Genome Haploid Length       365,165,005 bp   367,325,903 bp
Genome Repeat Length        213,881,262 bp   215,146,924 bp
Genome Unique Length        151,283,743 bp   152,178,979 bp
Model Fit                   57.1831%         93.7891%
Read Error Rate             0.318804%        0.318804%
```

##### SMUDGEPLOT

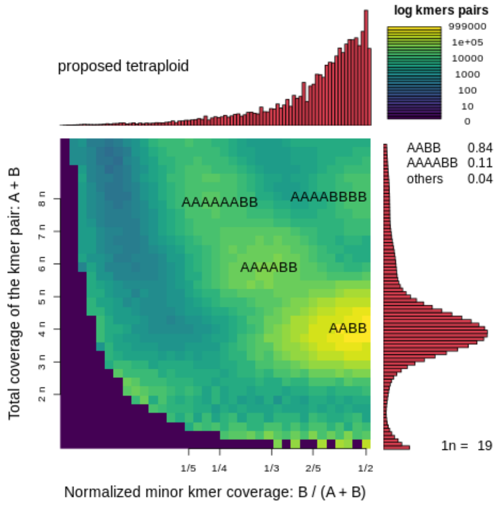

##### TETMER

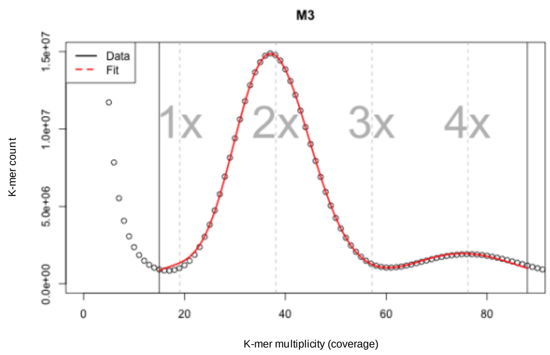

```
ALLOTETRAPLOID MODEL, AUTO FITTED
    haploid k-mer cov: 19.1
      per k-mer theta: 0.0198
                    T: 65.13
hapl non-rep GS (Mbp): 198.8
    bias (peak width): 0.5
     per k-mer diverg: 1.288

STARTING RANGES (MIN MAX)
    haploid k-mer cov: 15 21
log10 per k-mer theta: -3.85 0.6
                    T: 30 100
hapl non-rep GS (Mbp): 148 279
    bias (peak width): 0.1 3
              x range: 15 88
```

#### *Euphrasia* sp. (X1, tetraploid)

##### *GenomeScpe2.0*

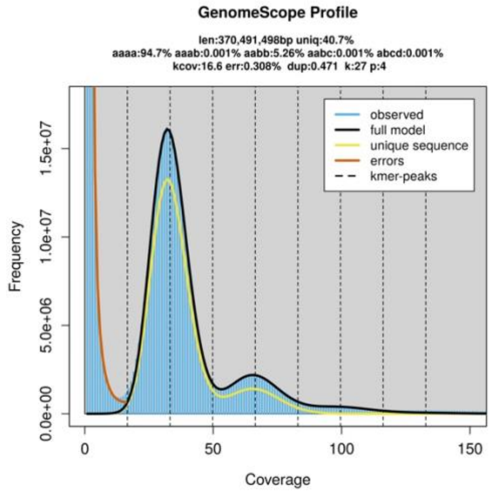

```
GenomeScope version 2.0
input file = /mnt/1TBSSD/Eukmers/numsE034
output directory = E034
p = 4
k = 27
initial kmercov estimate = 16

property                      min              max
Homozygous (aaaa)             92.6014%         94.9449%
Heterozygous (not aaaa)       5.05514%         7.39865%
aaab                          0%               0.79699%
aabb                          5.05514%         5.46933%
aabc                          0%               0.445348%
abcd                          0%               0.686974%
Genome Haploid Length         367,332,163 bp   370,491,498 bp
Genome Repeat Length          217,729,861 bp   219,602,503 bp
Genome Unique Length          149,602,302 bp   150,888,995 bp
Model Fit                     56.602%          94.8849%
Read Error Rate               0.307797%        0.307797%
```

##### *Smudgeplot*

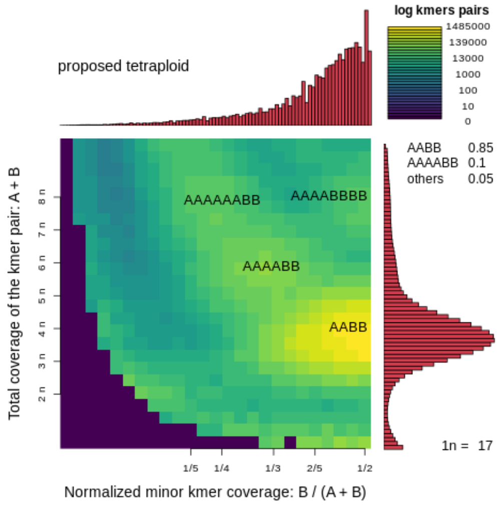

##### *Tetmer*

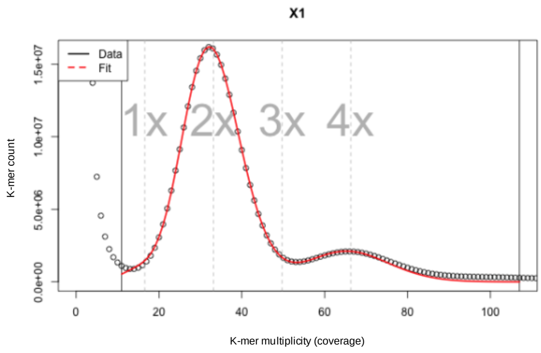

```
ALLOTETRAPLOID MODEL, AUTO FITTED
    haploid k-mer cov: 16.6
      per k-mer theta: 0.0158
                    T: 81.86
hapl non-rep GS (Mbp): 196
    bias (peak width): 0.5
     per k-mer diverg: 1.297

STARTING RANGES (MIN MAX)
    haploid k-mer cov: 15 19
log10 per k-mer theta: -4 -1.8
                    T: 0.001 100
hapl non-rep GS (Mbp): 20 235
    bias (peak width): 0.1 0.46
              x range: 11 107
```

#### *Euphrasia* sp. (X2, tetraploid)

##### GenomeScpe2.0

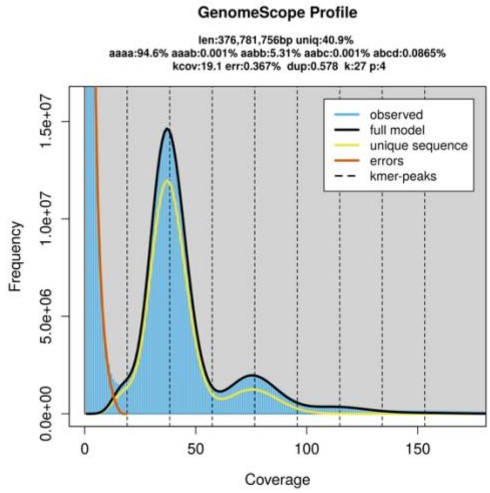

```
GenomeScope version 2.0
input file = /mnt/1TBSSD/Eukmers/numsE039
output directory = E039
p = 4
k = 27
initial kmercov estimate = 19

property                      min                max
Homozygous (aaaa)             92.6551%           94.9255%
Heterozygous (not aaaa)       5.0745%            7.3449%
aaab                          0%                 0.745196%
aabb                          5.0745%            5.54892%
aabc                          0%                 0.405733%
abcd                          0%                 0.645054%
Genome Haploid Length         374,819,298 bp     376,781,756 bp
Genome Repeat Length          221,669,233 bp     222,829,836 bp
Genome Unique Length          153,150,065 bp     153,951,920 bp
Model Fit                     56.9085%           94.3288%
Read Error Rate               0.367329%          0.367329%
```

##### Smudgeplot

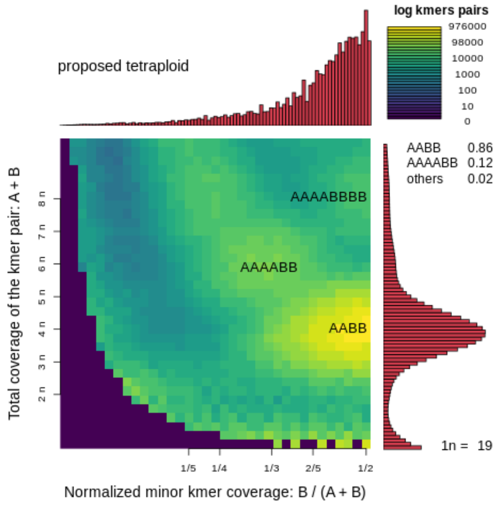

##### Tetmer

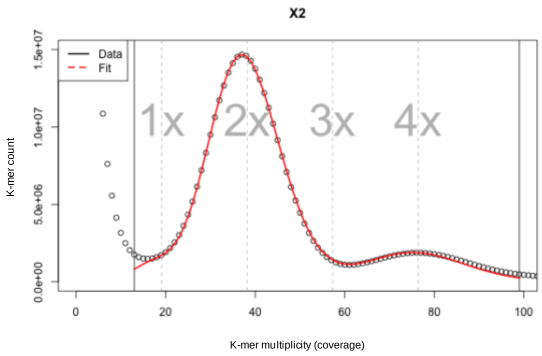

```
ALLOTETRAPLOID MODEL, AUTO FITTED
    haploid k-mer cov: 19.1
      per k-mer theta: 0.0259
                    T: 50.03
hapl non-rep GS (Mbp): 202.6
    bias (peak width): 0.6
     per k-mer diverg: 1.298

STARTING RANGES (MIN MAX)
    haploid k-mer cov: 13 20
log10 per k-mer theta: -2 0.6
                    T: 0.001 100
hapl non-rep GS (Mbp): 20 2000
    bias (peak width): 0.1 3
              x range: 13 99
```
